# Attracting pollinators vs escaping herbivores: eco-evolutionary dynamics of plants confronted with an ecological trade-off

**DOI:** 10.1101/2021.12.02.470900

**Authors:** Youssef Yacine, Nicolas Loeuille

## Abstract

Many plant traits are subject to an ecological trade-off between attracting pollinators and escaping herbivores. The interplay of both plant-animal interaction types determines their evolution. As most studies focus on either pollination or herbivory, how they jointly affect the eco-evolutionary dynamics of plant-animal communities is often left unknown. Within a plant-pollinator-herbivore community where interaction strengths depend on trait matching, we consider the evolution of a plant trait involved in both plant-animal interactions. Using adaptive dynamics, we uncover when stabilizing, runaway (i.e. directional) or disruptive selection emerges and its consequences for multispecies coexistence. We find that strong pollination relative to herbivory favors stabilizing selection and coexistence. Strong herbivory relative to pollination fosters runaway selection and threatens coexistence. Importantly, given balanced interactions, joint effects may lead to disruptive selection, allowing the emergence of plant dimorphism. The strength of the ecological trade-off largely explains the occurrence of these contrasting eco-evolutionary dynamics. In particular, plant diversification requires strong trade-offs, with the strongest trade-offs allowing long-term polymorphism. We discuss how our results relate to various empirical cases where the interplay of pollination and herbivory maintains plant polymorphism. Beyond maintenance, our work suggests that it might also have fueled the diversification process itself.

**Graphical Abstract:** Eco-evolutionary dynamics resulting from the evolution of plant phenotype under ecological trade-off
**A. Typical eco-evolutionary landscape**. The type of selection and the ecological outcome depend on the dissimilarity between animal phenotypes (i.e. preferences for plant phenotype), which is a proxy for the strength of the ecological trade-off. **B. The long-term community composition** depends on the type of selection. **(1)** Runaway selection leads to the extinction of a first animal species as the plant phenotype is diverging. **(a)** Pollinators are lost first so that runway selection continues until herbivores are also lost. **(b)** Herbivores are lost first so that selection turns stabilizing over time, leading to a perfect plant-pollinator matching. **(2)** Stabilizing selection can enable the maintenance of coexistence. **(3)** Disruptive selection leads to the emergence of plant dimorphism.

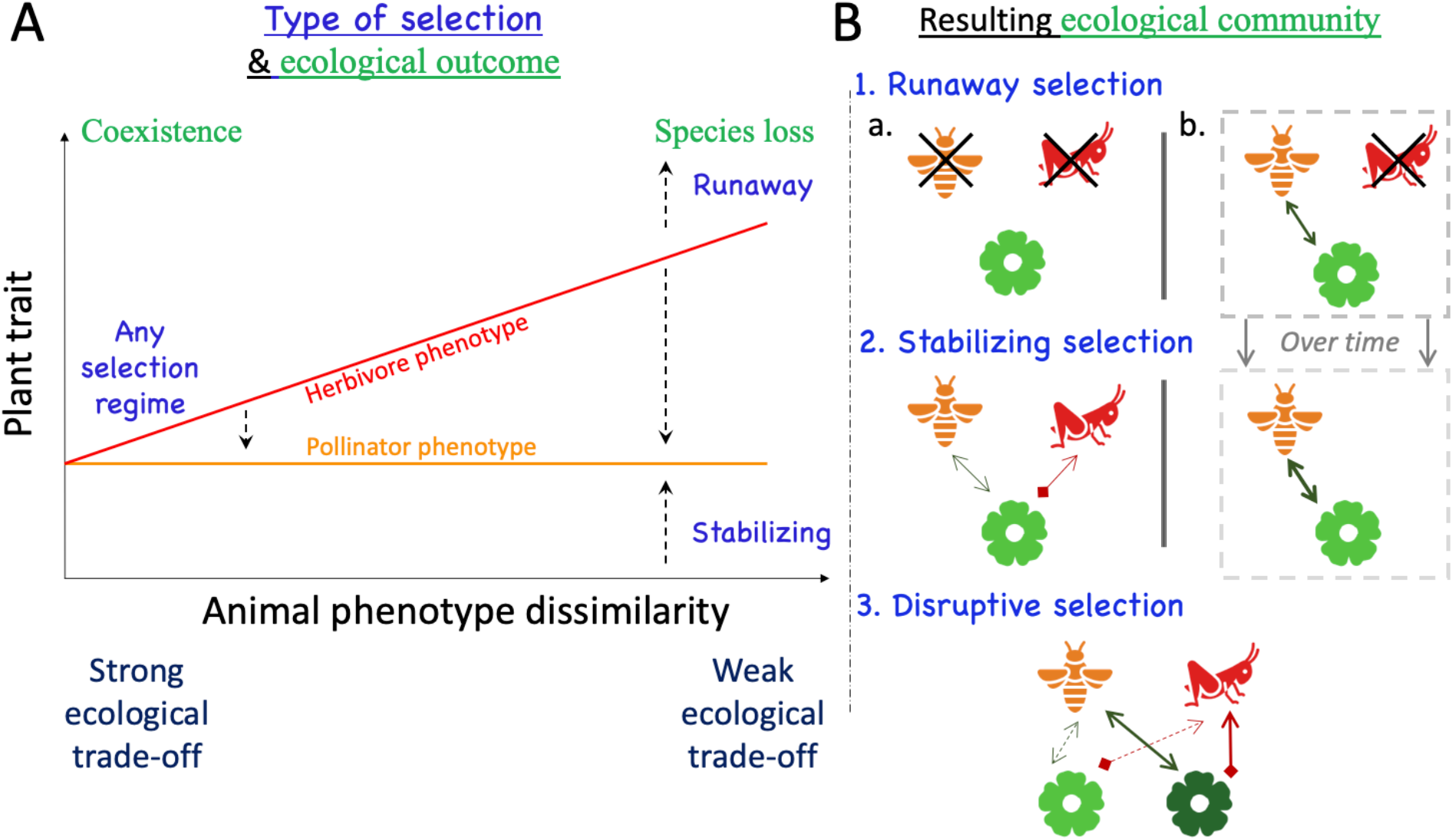

## Introduction

Flowering plants (i.e. angiosperms) are the most diverse and successful plant clade in terrestrial ecosystems, representing almost 90% of the species described (Hernández-Hernández & Wiens 2020). Most of them rely on animals for pollination (Ollerton *et al*. 2011). Because animal pollination can favor reproductive isolation, it has been proposed early as an important diversity driver among angiosperms (Grant 1949). Plant-herbivore interactions have, however, also been identified as potentially fostering diversification due to the ensuing evolutionary arms race between interacting antagonists (Ehrlich & Raven 1964). These two non-mutually exclusive hypotheses are supported by several phylogenetic investigations. Plant-pollinator coevolution explains higher diversification rates within angiosperms (Hodges & Arnold 1995; Sargent 2004; Hernández-Hernández & Wiens 2020), but plant-herbivore coevolution also spurs plant diversification as a result of defense-counterdefense innovations (Farrell *et al*. 1991; Becerra *et al*. 2009). Plant phylogenies reveal that plant-pollinator and plant-herbivore coevolution are often inextricably intertwined (e.g. Armbruster 1997), which advocates for an integrative framework accounting for both interaction types (Fontaine *et al*. 2011). From an ecological perspective, the positive or negative outcome of many interactions is often context-dependent (Chamberlain *et al*. 2014), varying for instance with the phenotype of interacting individuals (e.g. larvae vs. adult of nursery pollinators, Hahn and Brühl 2016). This dynamic nature of interactions along a mutualistic-antagonistic continuum (Thompson 1988; Gómez *et al*. 2023) further highlights the relevance of such an integrative framework.

A large number of plant traits are notably under conflicting selection due to the interplay between pollination and herbivory (Strauss & Irwin 2004), floral traits in particular (Strauss & Whittall 2006). Conflicting selection pressures can emerge from several mechanisms acting either in isolation or synergistically. Herbivory-induced changes in plant chemistry potentially reduce pollinator visitations (e.g. Kessler *et al*. 2011). Many plant traits acting on the plant visibility (size, phenology, floral display, volatile compounds) may increase herbivory (apparency hypothesis, Feeny 1976) while also attracting allies (e.g. Brody 2008). Genetic correlation can also exist between two plant traits, each involved in one plant-animal interaction (Strauss *et al*. 2004). A decisive consequence is that the selection pressures in the absence of either one animal species would be modified in the presence of both animal groups (Ramos & Schiestl 2019). Conflicting selection is very often due to shared preferences for plant phenotypes between pollinators and herbivores, a pattern that is largely widespread in nature (Strauss & Whittall 2006) and that promotes the stable coexistence of the community (Yacine & Loeuille 2022). This preference pattern indicates that plant species are subject to an ecological trade-off between attracting pollinators and escaping herbivores (Strauss *et al*. 2002). In other words, an increase in the strength of pollination (e.g. investment in attraction) comes at the cost of an increase in the strength of herbivory, while a decrease in the strength of herbivory (e.g. production of defenses) comes at the cost of a decrease in the strength of pollination. This trade-off has interestingly been shown to support the maintenance of a flower-color polymorphism in the wild radish *Raphanus sativus* (Irwin *et al*. 2003), which suggests that the interplay between pollination and herbivory could also lead to disruptive selection and promote diversification.

Conflicting selection arises because pollination and herbivory exert opposite pressures on plant traits (Thompson 2009), with contrasting implications in terms of diversification potential (Yoder & Nuismer 2010). These differences are especially prominent under the assumption that interaction strengths increase with the matching between plant and animal phenotypes (i.e. trait-matching). Such an assumption seems reasonable as it has been reported to apply in various instances, including flower color (Irwin *et al*. 2003) or flower display (Galen & Cuba 2001) matching animal preferences, chemical volatiles (Theis *et al*. 2014) matching animal detection abilities, or plant phenology matching that of animals (Brody 2008). As fitness increases with the strength of a mutualistic interaction, plants are selected to better match their pollinator phenotype and vice versa (e.g. spur length and pollinator tongue, Whittall & Hodges 2007) so that stabilizing selection is expected (e.g. Kiester *et al*. 1984). Pollination-induced stabilizing selection has been empirically characterized several times (e.g. Parachnowitsch and Kessler 2010; Sahli and Conner 2011; De Jager and Peakall 2019). It disfavors extreme phenotypes, thus reducing the potential for disruptive selection (Kopp & Gavrilets 2006; Yoder & Nuismer 2010; Maliet *et al*. 2020). Conversely, predation reduces prey fitness so that plants are selected to escape herbivore preferences (e.g. chemical defenses and herbivore tolerance, Becerra *et al*. 2009). Such arms race dynamics (Ehrlich & Raven 1964; Dawkins & Krebs 1979) expose the plant species to runaway selection, as observed in empirical systems (e.g. Mauricio and Rausher 1997; Becerra et al. 2009; Griese et al. 2021). Because apparent competition (Holt 1977) imposes a cost on phenotype matching between conspecific plants, herbivory also favors increased plant phenotypic divergence thereby enhancing disruptive selection (Kopp & Gavrilets 2006; Yoder & Nuismer 2010; Maliet *et al*. 2020). Note finally that conflicting selection may have far-reaching ecological implications. The relative interaction strength of pollination vs. herbivory has been shown to determine coexistence within plant-pollinator-herbivore communities (Mougi & Kondoh 2014; Sauve *et al*. 2016a; Yacine & Loeuille 2022). Because interaction strengths are the result of plant-animal coevolution, the evolution of plant traits under conflicting selection may have important consequences for multispecies coexistence.

In the present paper, our goals are to determine how the interplay of pollination and herbivory drives the evolution of plant traits involved in both interactions, and how, in turn, such an evolution affects the maintenance of coexistence. We model a community – a plant species interacting simultaneously with a pollinator and a herbivore species - in which plant-animal interactions depend on a plant trait involved in both interactions. Our previous purely ecological investigation of a similar three-species system has shown that a balance between the strength of plant-animal interactions makes the three-species coexistence more likely (Yacine & Loeuille 2022). Ignoring all evolutionary aspects, this previous study utterly focused on the ecological dynamics and their stability in relationship with plant-animal interaction strengths considered as independently varying parameters. In contrast, pollination and herbivory are here coupled by a focal plant trait whose phenotype determines the strength of both interactions (see Strauss & Whittall (2006) for examples). Each interaction increases in strength with the matching between plant phenotype and animal preferences (or more generally and henceforth, animal phenotypes). Under this trait-matching setting, we study the eco-evolutionary dynamics resulting from the evolution of the plant phenotype using the adaptive dynamics framework (Metz *et al*. 1992; Dieckmann & Law 1996).

We are particularly interested in understanding how these dynamics depend on an ecological trade-off to which the plant might be subject. An ecological trade-off is here defined as a positive covariation between the strengths of pollination and herbivory, i.e. stronger pollination (resp. weaker herbivory) comes at the cost of stronger herbivory (resp. weaker pollination). We first show that under our trait-matching assumption, variations in plant phenotype intrinsically entail an ecological trade-off whose strength depends on the dissimilarity between animal phenotypes (details in Methods, **Fig. 1**). Manipulating animal phenotype dissimilarity, we investigate how the strength of this ecological trade-off affects the type of selection on the plant trait and the maintenance of coexistence. When animal phenotypes are highly dissimilar (weak trade-off), interaction strengths are intrinsically unbalanced making the plant essentially interacting with one animal species so that we expect selection to be stabilizing close to the pollinator phenotype, and runaway close to the herbivore phenotype. Coexistence should then be frequently disrupted as stable coexistence requires the plant-animal interaction strengths to be of similar magnitudes (Yacine & Loeuille 2022). When animal phenotypes are fairly similar (strong trade-off), the balance between interaction strengths should, in contrast, favor the maintenance of coexistence. The potential for disruptive selection to occur in such instances, and the maintenance of the ensuing polymorphism, is one of the key questions of the present study.

**Fig. 1:**
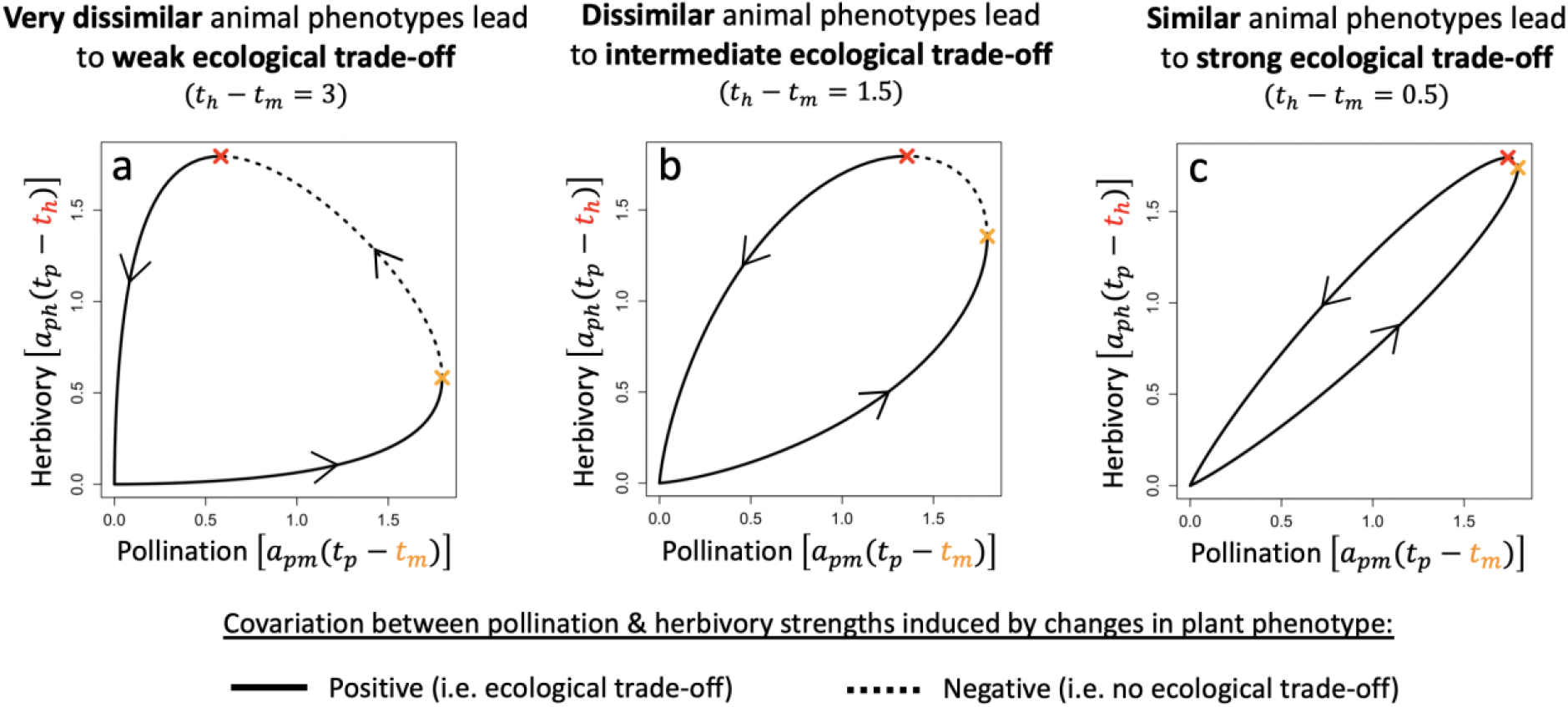
Animal phenotype dissimilarity determines the strength of the ecological trade-off. Each graph shows the covariation between the strength of plant-animal interactions resulting from varying plant phenotype (*t*_*p*_ varies from −10 to 10 in the direction indicated by the arrows, the orange (resp. red) point indicates when *t*_*p*_ = *t*_*m*_ (resp. *t*_*p*_ = *t*_*h*_)). **a. b. c. Increasing strength of ecological trade-off with increasing similarity between animal phenotypes (i.e. decreasing dissimilarity)**. <u>Parameter set:</u> ecological (*r*_*p*_ = 10, *r*_*m*_ = −1, *r*_*h*_ = −4, *c*_*p*_ = 0.6, *c*_*m*_ = 0.5, *c*_*h*_ = 0.4, *e*_*m*_ = 0.2, *e*_*h*_ = 0.3); interspecific (*t*_*m*_ = 0, *a*_*pm*0_ = *a*_*ph*0_ = 9, *σ*_*Pol*_ = *σ*_*Her*_ = 2).

## Model & Methods

### Model

#### Population dynamics

We model the dynamics of a plant-pollinator-herbivore module (*P*,*M*,*H* respectively) using a Lotka-Volterra framework:

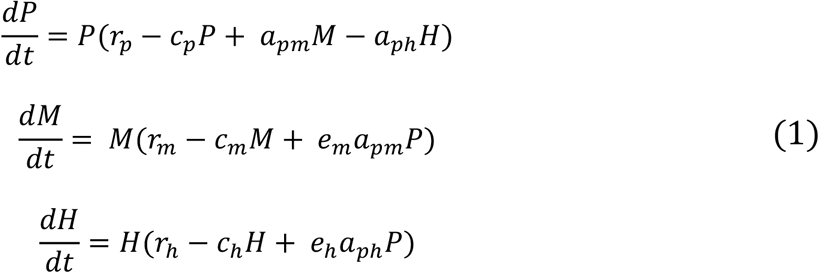

Plants are assumed to have a positive intrinsic growth rate (*r*_*p*_ > 0), while both pollinator (*r*_*m*_ < 0) and herbivore growth rates (*r*_*h*_ < 0) are assumed negative. As in previous models (e.g. Sauve et al. 2014), we thus assume the plant-animal interaction to be obligate for the animals and facultative for the plant. All interacting species undergo negative (quadratic) density-dependence. These terms are here interpreted as intraspecific competition. The plant competition rate (*c*_*p*_) would notably encompass the competition for space, water or nutrients (Craine & Dybzinski 2013). Animal competition rates (*c*_*m*_, *c*_*h*_) would essentially coincide with interference (e.g. Thébault and Fontaine 2010). These terms might however be more generally considered as any negative regulation not explicitly accounted for in our model (e.g. other predators or resources, diseases….). The per-capita strengths of interspecific interactions are given by *a*_*pm*_ for pollination and *a*_*ph*_ for herbivory. *e*_*m*_ and *e*_*h*_ are the conversion efficiencies from plants to animals. Table 1 recapitulates model variables and parameters. Our formulation of population dynamics implies an indirect (i.e. plant-mediated) interaction between animal densities, but no direct interaction as would for instance be the case when an animal behaves as both pollinator and herbivore (e.g. nursery pollinators).

**Table 1:** List of all model parameters and variables with their biological definitions, values and dimensions.

| Variables and parameters |  | Biological definition | Value | Dimension |
| --- | --- | --- | --- | --- |
| Variables | $P$ | Plant biomass density | | $mass.length^{-2}$ |
| | $M$ | Pollinator biomass density | | $mass.length^{-2}$ |
| | $H$ | Herbivore biomass density | | $mass.length^{-2}$ |
| | $t_p$ | Plant phenotype | | *Dimensionless* |
| *Animal phenotypes* | $t_m$ | Pollinator phenotype | 0 | *Dimensionless* |
| | $t_h$ | Herbivore phenotype | [0,3] | *Dimensionless* |
| *Interspecific interaction parameters* | $t_h - t_m$ | Animal phenotype dissimilarity<br>Proxy for the strength of the ecological trade-off (see Methods) | [0,3] | *Dimensionless* |
| | $a_{pm0}$ | Basal pollination rate (per capita) | [3,9] | $(M.L^{-2})^{-1}.t^{-1}$ |
| | $a_{ph0}$ | Basal herbivory rate (per capita) | [3,9] | $(M.L^{-2})^{-1}.t^{-1}$ |
| | $\sigma_{Pol}$ | Pollination niche width | [1.5,3] | *Dimensionless* |
| | $\sigma_{Her}$ | Herbivory niche width | [1.5,3] | *Dimensionless* |
| *Other ecological parameters* | $r_p$ | Plant intrinsic growth rate | 10 | $time^{-1}$ |
| | $r_m$ | Pollinator intrinsic growth rate | -1 | $time^{-1}$ |
| | $r_h$ | Herbivore intrinsic growth rate | -4<br>(-1, -1) | $time^{-1}$ |
| | $c_p$ | Plant intra-specific competition rate | 0.6<br>(0.6, 1.7) | $[(mass.length^{-2}).time]^{-1}$ |
| | $c_m$ | Pollinator intra-specific competition rate | 0.5<br>(0.4, 0.7) | $[(mass.length^{-2}).time]^{-1}$ |
| | $c_h$ | Herbivore intra-specific competition rate | 0.4 | $(M.L^{-2})^{-1}.t^{-1}$ |
| | $e_m$ | Plant to pollinator conversion efficiency | 0.2 | *Dimensionless* |
| | $e_h$ | Plant to herbivore conversion efficiency | 0.3<br>(0.2, 0.3) | *Dimensionless* |
| *Numerical simulations* | $\mu$ | Mutation probability (per unit of time and plant biomass density) | $2.10^{-7}$ | $[(mass.length^{-2}).time]^{-1}$ |
| | $\sigma$ | Mutation amplitude (standard deviation) | 0.02 | *Dimensionless* |
| | $\varepsilon$ | Extinction threshold | $2.10^{-6}$ | $mass.length^{-2}$ |

column “Value”, “*other ecological parameters*”: in parenthesis are provided the values for the robustness ecological parameter sets when they differ from that of our main ecological parameter set (see section Methods/Workflow).

Without evolution, stable three-species coexistence occurs when the strengths of pollination (*a*_*pm*_) and herbivory (*a*_*ph*_) are balanced (Yacine & Loeuille 2022). When pollination is much stronger than herbivory, stability may be lost in the form of unbounded population growth (e.g. Fig. B1, Fig. 5b-d). In this parameter region (*a*_*pm*_ *≫ a*_*ph*_), our model thus fails at producing biologically realistic dynamics, which indicates that other ignored processes are prominent in such instances (e.g. saturating parameters, wider community context; discussed extensively in Yacine & Loeuille, 2022). One of our main previous results, however, is that the plant-herbivore interaction largely reduces the size of this parameter region, i.e. herbivory often stabilizes unbounded dynamics in plant-insect systems (see Fig. B1.A in appendix B). In the present work, we investigate how the relative strength of pollination vs. herbivory influences plant evolution, and how such evolution, in turn, affects the maintenance of multispecies coexistence. Evolution is characterized in the parameter region over which stable three-species coexistence is obtained (analytical expression of equilibrium (*P*^*^, *M*^*^, *H*^*^) given in Appendix A.I, Supporting Information), as well as in regions of unbounded growth (see Methods).

#### Plant-animal interactions depend on trait matching

We assume plant-animal interactions to intensify as the matching between plant (*t*_*p*_) and animal phenotypes – pollinator (*t*_*m*_) or herbivore (*t*_*h*_) – increases (equations 2). Interactions are maximal when traits perfectly match (i.e. |*t*_*p*_ − *t*_*m*_| = 0, resp. |*t*_*p*_ − *t*_*h*_| = 0), and weaken when trait-matching is reduced (i.e. |*t*_*p*_ − *t*_*m*_| ↗, resp. |*t*_*p*_ − *t*_*h*_| ↗). Examples include the color of plant flowers (*t*_*p*_) and associated animal preferences (*t*_*m*_, *t*_*h*_) (e.g. Irwin *et al*. 2003) or phenological traits (e.g. Brody 2008) such as the date of flowering (*t*_*p*_) and of animal activity (*t*_*m*_, *t*_*h*_).

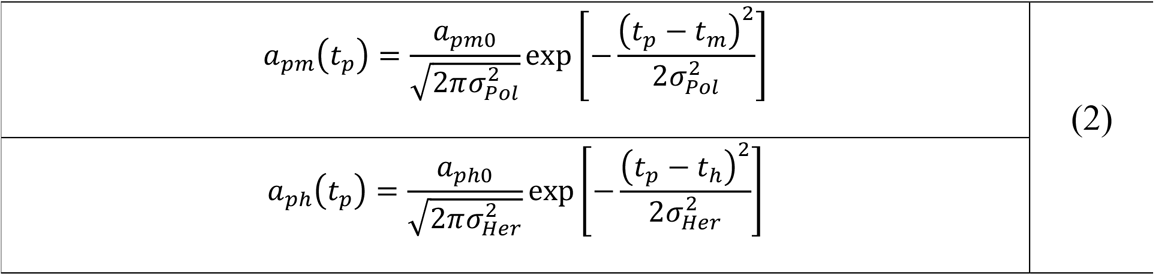

As *σ*_*Pol*_ controls how quickly the strength of pollination decreases with plant-pollinator phenotype dissimilarity, it corresponds to the pollination-niche width, which depends on the generalism of both species. Likewise, *σ*_*Her*_ embodies the herbivory-niche width. *a*_*pm*0_ and *a*_*ph*0_ correspond to basal interaction rates. See table 1 for details.

## Methods

### Definition and emergence of an ecological trade-off

An ecological trade-off between attracting pollinators and escaping herbivory is defined as a positive covariation between the strengths of pollination and herbivory, i.e. both increasing or decreasing. As our focal plant trait is involved in both plant-animal interactions (equation 2), its variation affects both interaction strengths (e.g. **Fig. 1b**). Because of our trait-matching hypothesis, the induced covariation between the strengths of pollination and herbivory is positive when the plant phenotype is outside the phenotypic interval [*t*_*m*_, *t*_*h*_] *V*ariation in plant phenotype there entails an ecological trade-off (**Fig. 1b, solid line**). Within the interval [*t*_*m*_, *t*_*h*_], there is no ecological trade-off (**Fig. 1b, dotted line**).

We consider the dissimilarity between animal phenotypes (*t*_*h*_ − *t*_*m*_) to be a proxy for the strength of the ecological trade-off that emerges within our framework. There are two reasons for that. First, as animal phenotype similarity increases (**Fig. 1, a vs. b vs. c**), the phenotypic region over which plant phenotype variation induces an ecological trade-off increases in size. Second, over this region, in the case of very dissimilar animal phenotypes (i.e. *t*_*m*_ ≪ *t*_*h*_, **Fig. 1a**), plant phenotype variations affect much more one interaction than the other, depending on the closest animal phenotype. In contrast, over the same region but in the case of similar phenotypes (i.e. *t*_*m*_ ≈ *t*_*h*_, **Fig. 1c**), any variation of the plant phenotype has an effect of comparable magnitude on the strength of both interactions. All in all, the ecological trade-off between attracting pollinators and escaping herbivores gets stronger as animal dissimilarity decreases (**Fig. 1**).

### Adaptive dynamics and type of selection

Within a monomorphic plant population with phenotype *t*_*p*_ (resident), we investigate whether a mutant with a new phenotype *t*_*p*_′ can invade. Invasion fitness *w*(*t*_*p*_′, *t*_*p*_) is computed as the per capita growth rate of that mutant, when rare and in the environment (*P*^*^, *M*^*^, *H*^*^) set by the resident population (Metz *et al*. 1992, see appendix A.II). When a mutant invades (i.e. *w*(*t*_*p*_′, *t*_*p*_) > 0), it replaces the resident population, thus becoming the new resident. The sequence of trait substitutions defines the long-term evolutionary dynamics. Assuming small and rare mutational steps, the variation of phenotype *t*_*p*_ is proportional to the selection gradient (equation (3)), i.e. the derivative of invasion fitness with respect to the mutant’s trait (Dieckmann & Law 1996). Evolutionary singularities 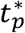 correspond to phenotypes that nullify the selection gradient.

The type of selection – stabilizing, disruptive, or runaway – acting on the plant trait depends on (1) the properties of the evolutionary singularities, and (2) the position of the plant phenotype relative to these singularities. Two independent properties - convergence and invasibility – characterize evolutionary singularities (criteria in appendix A.II). Convergence indicates whether the trait evolves toward the singularity in its vicinity. Two types of singularities are convergent – the non-invasible continuously stable strategy (CSS, Eshel, 1983) and the invasible branching point (Geritz *et al*. 1997). Invasibility specifies whether the singularity may be invaded by nearby mutants (i.e. ESS, Maynard Smith & Price 1973). CSS phenotypes correspond to cases of stabilizing selection (e.g. Fig. 2B.a). Selection is thus considered stabilizing in the basin of attraction of a CSS. Plant-pollinator-herbivore coexistence is notably maintained if a CSS is reached while the three species coexist. We also describe directional selection towards and within a phenotypic region in which our model produces unbounded growth as stabilizing selection (in terms of purely evolutionary dynamics, not ecological). This choice was motivated by both mathematical and biological coherence (details in appendix B.II.1). Unbounded growth regions are notably attractive in terms of evolution (i.e. convergence, note that classical tools of adaptive dynamics (e.g. selective gradient) do not apply within these regions but only in their vicinity). Our choice also preserves the association between stabilizing selection and coexistence maintenance as areas of unbounded growth are regions of phenotypic space in which our model fails to produce realistic dynamics, but in which, from a biological point of view, coexistence should be maintained (notion of “permanent coexistence”, Hutson and Law 1985, discussed in Yacine and Loeuille 2022). In contrast to a CSS, a branching point yields the emergence of plant dimorphism due to disruptive selection (e.g. Fig. 2B.d). Accordingly, selection is considered disruptive in the basin of attraction of a branching point. Finally, phenotypes that are not in the basin of attraction of a convergent singularity are under runaway selection. This is possible in the presence of a non-convergent singularity, i.e. a repellor. For the sake of simplicity, we will refer to this set of phenotypes as the basin of attraction associated with runaway selection. Runaway selection should always disrupt plant-pollinator-herbivore coexistence (e.g. Fig. 2B.b). We illustrate how the proportion of phenotypic space under each type of selection is calculated in Appendix B.II.1.

**Fig. 2:**
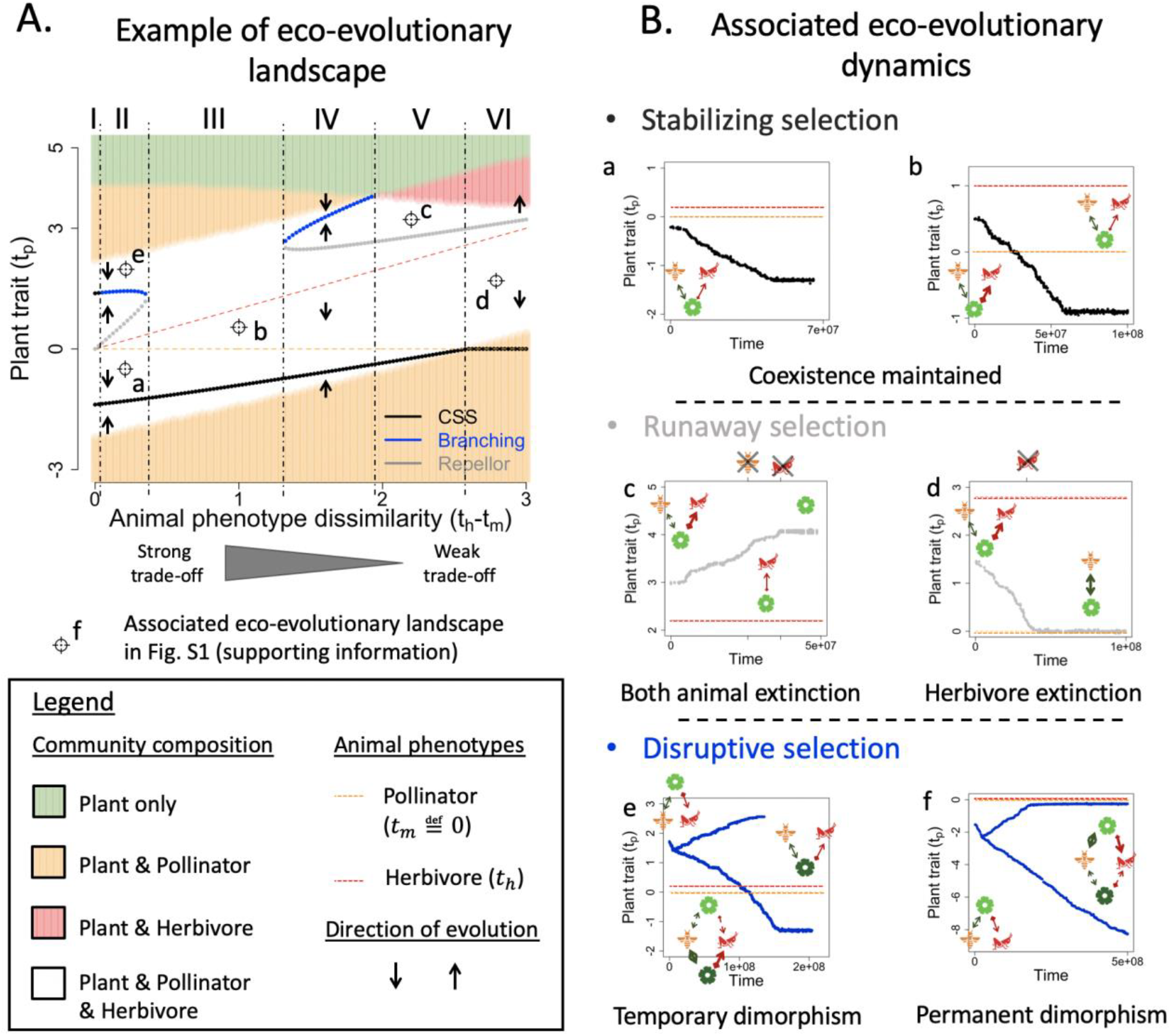
Typical examples of eco-evolutionary dynamics with their associated eco-evolutionary landscape. **A. Eco-evolutionary landscape**. The points labelled by letters indicate the initial conditions of the simulations presented in B. <u>Ecological parameter set:</u> (*r*_*p*_ = 10, *r*_*m*_ = −1, *r*_*h*_ = −4, *c*_*p*_ = 0.6, *c*_*m*_ = 0.5, *c*_*h*_ = 0.4, *e*_*m*_ = 0.2, *e*_*h*_ = 0.3). <u>Interspecific parameter set</u> (*a*_*pm*0_ = 5, *a*_*ph*0_ = 7, *σ*_*Pol*_ = 3, *σ*_*Her*_ = 2.8) **B. Simulated eco-evolutionary dynamics associated with each type of selection. a-b. Stabilizing selection enables the maintenance of coexistence** (*t*_*h*_ = 0.2 & 1 respectively). **c-d. Runaway selection leads to coexistence loss** (*t*_*h*_ = 2.8 & 2.2 respectively). **e-f. Disruptive selection allows the emergence of plant dimorphism** (*t*_*h*_ = 0.2 & 0.1 respectively). Note that for f, the interspecific parameter set is modified (*a*_*pm*0_ = 5, *a*_*ph*0_ = 9, *σ*_*Pol*_ = 1.7, *σ*_*Her*_ = 2). The associated landscape is given in **Fig. S1**. In B, pictograms depict the community composition, with arrow thickness correlating to interaction strengths.

### Numerical simulations of community eco-evolutionary dynamics

This framework is completed by numerical simulations starting from a monomorphic plant population (*t*_*p*_) interacting with a pollinator (*t*_*m*_) and a herbivore (*t*_*h*_) population. Mutations are randomly generated following a Poisson process characterized by a mutation probability per unit of time and plant biomass density *µ* = 2. 10^−7^. Proportionally to phenotype abundances, a parent phenotype is randomly chosen at each mutation event. The mutant phenotype is drawn from a Gaussian distribution centered around the parent phenotype with standard deviation *σ* = 0.02. Its initial density is set to a small value *ε*, taken from the parent population. Symmetrically, populations falling below *ε* are removed from the system (extinction).

### Workflow

Animal phenotypes are fixed parameters (without loss of generality: *t*_*m*_ = 0, *t*_*h*_ ≥ *t*_*m*_) while we study the evolution of the plant phenotype (*t*_*p*_). Parameters directly affecting plant-animal interactions – i.e. the interspecific parameter set (*t*_*h*_ − *t*_*m*_, *a*_*pm*0_, *a*_*ph*0_, *σ*_*Pol*_, *σ*_*Her*_) – are at the core of our investigation. Thanks to the model simplicity, several analytical results uncovering various aspects of evolutionary dynamics (e.g. equation (3)) are possible. For aspects that cannot be mathematically investigated, we provide numerical resolutions characterizing the variation of eco-evolutionary dynamics along the gradient of trade-off intensity (E3-diagrams, e.g. Fig. 2A). To broaden our understanding of possible evolutionary dynamics, we undertake Monte Carlo experiments (details in appendix B.II). We let interspecific parameters vary independently within their interval ranges (table 1), which were chosen to explore a wide range of pollination and herbivory intensities (Appendix B.I). Remaining parameters (*r*_*p*_, *r*_*m*_, *r*_*h*_, *c*_*p*_, *c*_*m*_, *c*_*h*_, *e*_*m*_, *e*_*h*_) are fixed, but the experiments were conducted for three different sets to assess the robustness of our results: a main ecological parameter set allowing a large range of plant-animal interaction strengths to be compatible with stable coexistence, a second one characterized by symmetrical parameter values for animals, and a third one in which unbounded population growth is made impossible (details in Appendix B.I). We highlight here that our results are essentially robust, as detailed hereinafter. The different Monte Carlo experiments are presented directly in the Result section when needed. Our approach is finally complemented by numerical simulations of the community eco-evolutionary dynamics. Such simulations are used to investigate the emergence of plant dimorphism in the case of disruptive selection.

## Results

### A typical example of eco-evolutionary dynamics

Eco-evolutionary dynamics qualitatively depend on the strength of the ecological trade-off (Fig. 2). These dynamics are characterized by stabilizing or disruptive selection when trade-offs are strong. They then allow the coexistence of all species. Such a coexistence is, however, often lost in the case of weak trade-offs, where runaway selection is much more common. As developed later on (Fig. 4A), these variations are more general than the specific example we introduce here (Fig. 2A). On Fig. 2A, the plant phenotype always converges towards a CSS phenotype under very strong trade-offs (region I) so that selection is stabilizing. As the trade-off weakens (region II), both stabilizing (Fig. 2B.a) and disruptive selection (Fig. 2B.e) become possible depending on the initial plant phenotype. Stabilizing (resp. disruptive) selection is observed over the basin of attraction of the CSS (black line) (resp. branching point, blue line). These basins of attraction (see arrows) are separated by the repellor (grey line). At weaker trade-offs (region III), the basin of attraction of the CSS covers the whole coexistence area (white) so that selection is always stabilizing (Fig. 2B.b). In region IV, eco-evolutionary dynamics are qualitatively similar to those of region II. For even weaker trade-offs (region V), runaway selection is observed whenever the initial plant phenotype is above the repellor (Fig. 2B.c). Associated eco-evolutionary dynamics lead first to the extinction of pollinators, then of herbivores. Below the repellor, selection stabilizes at the CSS phenotype. For even weaker trade-offs (region VI), the CSS exits the coexistence area so that only runaway selection remains possible. It provokes the extinction of both animal species when starting above the repellor, and that of herbivores when starting below the repellor. The extinction of herbivores then leads to a perfectly matched plant-pollinator community (Fig. 2B.d).

The ecological dynamics induced by each type of selection qualitatively differ (Fig. 2B). Coexistence is always maintained in the case of stabilizing selection. It is always disrupted in the case of runaway selection due to the weakening of one or both plant-animal interactions. Disruptive selection allows the emergence of plant dimorphism. The two plant phenotypes diverge, one leading to stronger plant-animal interactions, the other to weaker plant-animal interactions. Dimorphism can be temporary (Fig. 2B.e) or permanently maintained (Fig. 2B.f). An example of eco-evolutionary landscape associated with permanent dimorphism is given in Fig. S1 (Supporting Information), while the landscape presented in Fig. 2A typically leads to temporary branchings. In addition to trade-off intensity, other interspecific parameters can thus be responsible for qualitative changes in eco-evolutionary dynamics. Most of such (qualitative) variation is covered by the simulations in Fig. 2B.

### Opposite effects of pollination vs herbivory on the (local) selection gradient

The selection gradient (equation (3)) shows that plant evolution is under two contrasting selective forces (i.e. opposite sign).

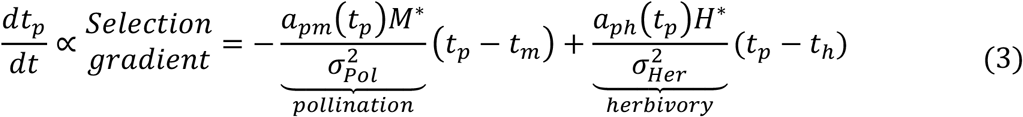

Pollination selects for plant phenotypes that are closer to that of pollinators (negative sign). Herbivory selects for phenotypes that are further away from that of herbivores (positive sign). These two selective forces act synergistically when the plant phenotype is within [*t*_*m*_, *t*_*h*_] so that evolution enables a simultaneous increase in pollination and decrease in herbivory. In contrast, these two forces become antagonistic when the plant phenotype is outside [*t*_*m*_, *t*_*h*_]. Pollination selects for stronger plant-animal trait matchings, while herbivory selects for weaker trait matchings. Such conflicting selection captures the ecological trade-off with which plants are confronted.

Pollination and herbivory have also an opposite effect on the evolutionary stability – i.e. non-invasibility – of evolutionary singularities 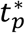. Singularities are invasible if inequality (4) is satisfied (proof and expression of 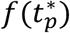 in Appendix A.II.2).

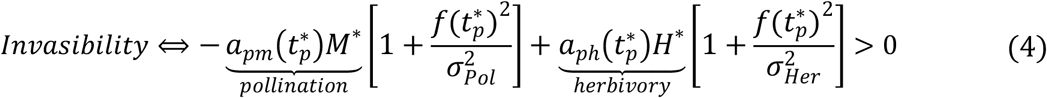

Invasibility is thus favored by herbivory and disfavored by pollination. Convergence being however not analytically tractable, we rely on a first Monte Carlo experiment (MC1, details in appendix B.II.2) to investigate the relationship between selection type and plant-animal interactions. We sampled 10 000 interspecific parameter sets and analyzed the nature of all evolutionary singularities allowing ecological coexistence, as well as interaction strengths and animal densities at these singularities. The resulting dataset thereby links the ratio of pollination to herbivory at the singularity 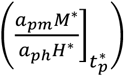 to the type of selection. Stabilizing, runaway and disruptive selections are characterized by contrasting pollination to herbivory ratios (**Fig. 3**), whose variation explains around two-thirds of the variance in selection type (Kruskal-Wallis).

**Fig. 3:**
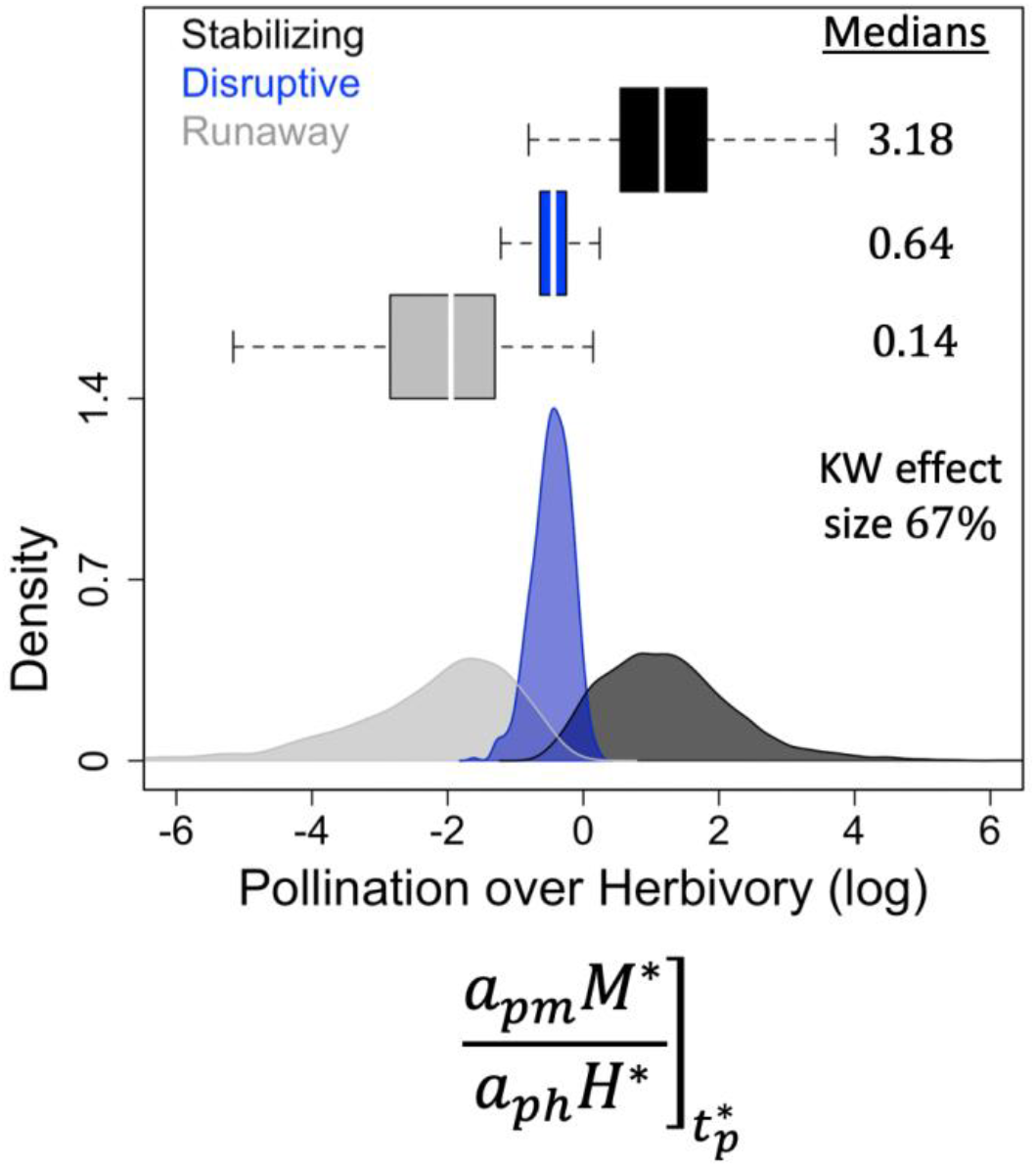
Distribution of pollination to herbivory ratios depending on the type of selection. The ratio is measured at the corresponding evolutionary singularity. The medians indicated correspond to absolute values (no log). The effect size of the Kruskal-Wallis test (*p*_*value*_ < 2.2 10^−16^) is also indicated. <u>Ecological parameter set:</u> (*r*_*p*_ = 10, *r*_*m*_ = −1, *r*_*h*_ = −4, *c*_*p*_ = 0.6, *c*_*m*_ = 0.5, *c*_*h*_ = 0.4, *e*_*m*_ = 0.2, *e*_*h*_ = 0.3). Results are from our first Monte Carlo experiment (MC1, appendix B.II.2), with 10 000 interspecific parameter sets sampled.

High pollination relative to herbivory is associated with stabilizing selection (Fig. 3). High herbivory relative to pollination is associated with runaway selection (Fig. 3). When pollination and herbivory are balanced, selection turns disruptive (Fig. 3). The association of stabilizing, runaway and disruptive selection with respectively high, low and balanced pollination to herbivory ratios implies that the convergence of evolutionary singularities (i.e. convergence stability, Eshel 1983) is favored by pollination and disfavored by herbivory. Disruptive selection requires a balance between pollination and herbivory because strong herbivory relative to pollination fosters invasibility (inequality (4)), but convergence is lost when herbivory is too strong, leading to runaway dynamics (Fig. 3). Note finally that simpler ratios measuring the strength of pollination vs. herbivory (e.g. interaction strengths 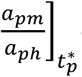 or animal densities 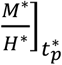) also have a large power in explaining the nature of selection (table S1, Supporting Material). These key results derived from our Monte Carlo experiment are robust to the modification of the ecological parameter set (Appendix C.I).

### Effects of interspecific parameters on selective landscapes

Beyond local selection at evolutionary singularities, we now investigate the effect of plant-animal interactions on the selective landscape at large, i.e. the distribution of basins of attraction associated with each type of selection (e.g. Fig. 2A). We build a second Monte Carlo experiment (MC2, details in Appendix B.II.3). For each focal interspecific parameter, we divide the potential range of that parameter (table 1) into six equal intervals to sample from, the sampling of the other interspecific parameters remaining unconstrained. For each sampling (6 x 1000), we calculate the proportion of phenotypic space associated with each type of selection (i.e. basin of attraction, see Appendix B.II.1 for details). We thereby estimate the effect of varying (1) the strength of ecological trade-off (Fig. 4A) and (2) the intensity and generalism of pollination and herbivory (Fig. 4B) on the selective landscape.

**Fig. 4:**
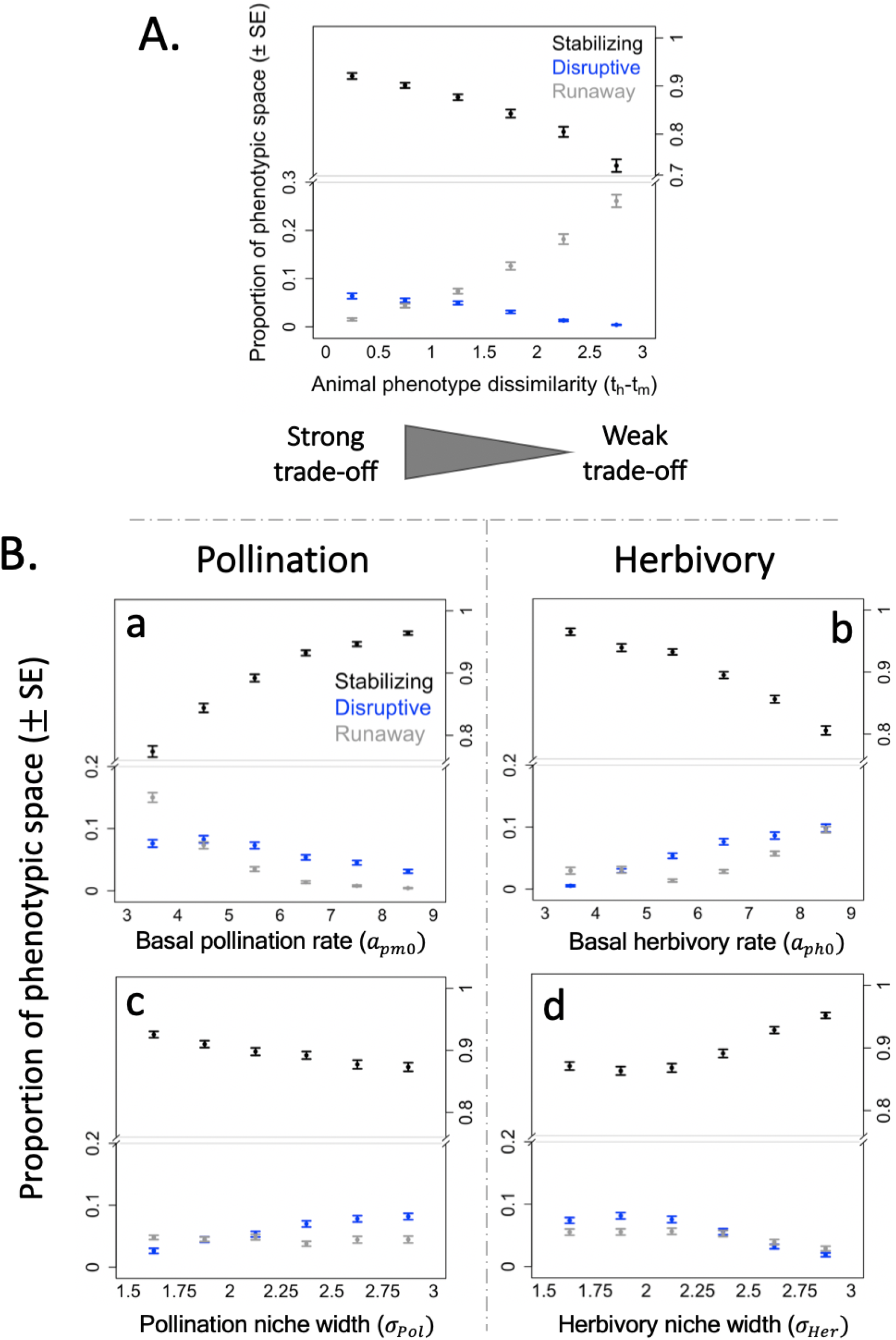
Effect of interspecific parameters on the selective landscape. **A. The selective landscape depends on trade-off intensity. B. Opposite effects of pollination and herbivory on selection**. Please note that animal phenotype dissimilarity (*t*_*h*_ − *t*_*m*_) was here further constrained within [0, 1.5]. This was done to better capture the effects of the other interspecific parameters on disruptive selection, the latter being hardly observed for high animal phenotype dissimilarities (i.e. for weak trade-offs, see Fig. 4A and main text). **a. b. Variations in basal interaction rates. c. d. Variations in niche widths**. Results are from our second Monte Carlo experiment (MC 2, appendix B.II.3), with 1000 interspecific parameter sets sampled at each point. Y-axes indicate the normalized size of the basins of attraction (Mean ± SE) associated with each type of selection (see appendix B.II.1). <u>Ecological parameter set:</u> (*r*_*p*_ = 10, *r*_*m*_ = −1, *r*_*h*_ = −4, *c*_*p*_ = 0.6, *c*_*m*_ = 0.5, *c*_*h*_ = 0.4, *e*_*m*_ = 0.2, *e*_*h*_ = 0.3). See Fig. C2 & C3 (Appendix C.II) for robustness.

Stabilizing selection dominates the selective landscape (Fig. 4), indicating that evolutionary dynamics usually tend to maintain plant-animal coexistence. This result is essentially robust to the modification of the ecological parameter set (Appendix C.II, but see Fig. C3.B).

Disruptive selection is possible at strong trade-offs, but not at weak trade-offs that typically lead to stabilizing or runaway selection (Fig 4A). Disruptive selection indeed requires a balance between pollination and herbivory (Fig. 3). Such a balance has to be achieved over a phenotypic region outside [*t*_*m*_, *t*_*h*_] for a branching point to occur. Phenotypes within [*t*_*m*_, *t*_*h*_] are continuously selected toward the pollinator phenotype and away from the herbivore one so that no singularity is possible between the two animal traits (see equation 3). When the trade-off is strong, balanced interactions naturally emerge as animal phenotypes are similar. When animals have very dissimilar phenotypes, pollination to herbivory ratios are either large or small outside [*t*_*m*_, *t*_*h*_], depending on which animal phenotype is closer to the plant phenotype. As a result, selection is either stabilizing or runaway at weak trade-offs. Note finally that disruptive selection is seldom observed with both our robustness ecological parameter sets (Fig. C2.A & C3.A, Appendix C.II), and that it similarly does not occur at weak trade-offs.

The reported opposite influence of pollination and herbivory on local selection extends in a consistent manner to the selective landscape (Fig. 4B). More precisely, the distribution of basins of attraction associated with each type of selection shifts towards a higher prevalence of stabilizing selection with stronger pollination, and a higher prevalence of runaway selection with stronger herbivory.

Increasing the basal rate of pollination (*a*_*pm*0_) favors stabilizing selection at the expense of disruptive and runaway selection (Fig. 4B.a). On the contrary, higher herbivory rates (*a*_*ph*0_) favor disruptive and runaway selection at the expense of stabilizing selection (Fig. 4B.b). These results suggest that the prevalence of disruptive selection is restricted by strong pollination and fostered by strong herbivory (Fig 4B. a vs. b). A narrower pollination niche width (*σ*_*Pol*_) increases the prevalence of stabilizing selection at the expense of disruptive selection (Fig. 4B.c). In contrast, disruptive and runaway selection become more frequent as the herbivory niche width (*σ*_*Her*_) gets narrower (Fig. 4B.d). Variations are, however, less pronounced for niche widths (*σ*) than for interaction rates (*a*_0_). Moreover, only in the case of interaction rates are these patterns robust to the variation of the ecological parameter set (Appendix C.II). While basal rate variations have a consistent effect on interaction strength across the phenotypic space, niche width variations increase or decrease interaction strength depending on plant-animal trait matching (i.e. *t*_*p*_ − *t*_*m*_, *t*_*p*_ − *t*_*h*_), likely explaining such a difference.

Changes in the niches of pollination and herbivory modify community eco-evolutionary dynamics, but these modifications are mainly driven by changes in evolutionary dynamics at strong trade-offs, and changes in ecological dynamics at weak trade-offs (Fig. 5 & S2, Supporting Material).

**Fig. 5:**
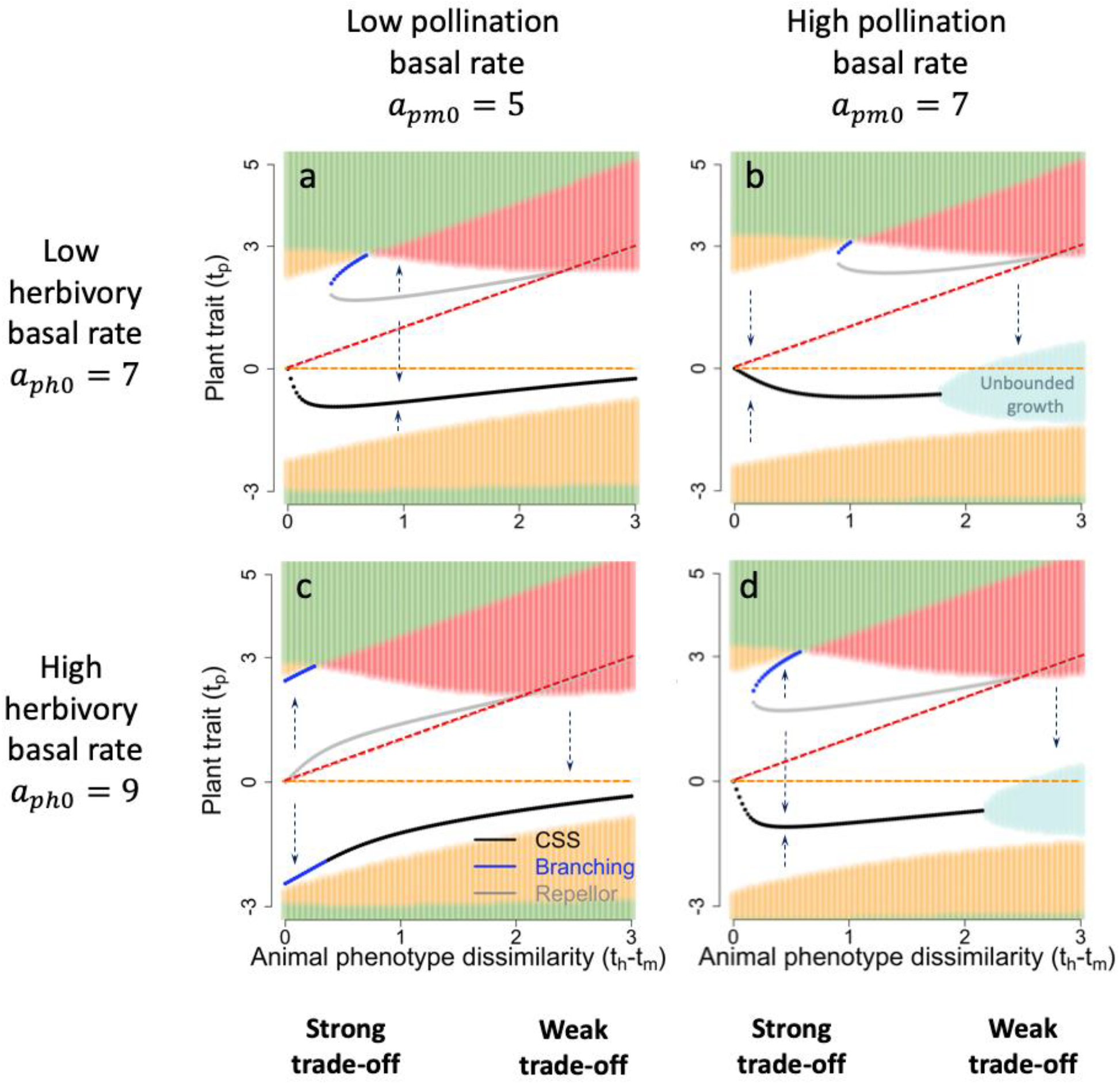
Effect of the basal rates of plant-animal interactions on community eco-evolutionary dynamics. At strong trade-offs, eco-evolutionary dynamics are primarily altered by changes affecting evolutionary singularities. At weak trade-offs, the alteration is mainly mediated by changes in the composition of the ecological community (i.e. background colors). Orange and red dotted lines indicate the pollinator and herbivore phenotype. Arrows indicate evolutionary trajectories and background colors indicate community composition as in Fig. 2A, with light blue figuring regions in which our model fails to produce realistic dynamics (unbounded population growth). <u>Ecological parameter set:</u> (*r*_*p*_ = 10, *r*_*m*_ = −1, *r*_*h*_ = −4, *c*_*p*_ = 0.6, *c*_*m*_ = 0.5, *c*_*h*_ = 0.4, *e*_*m*_ = 0.2, *e*_*h*_ = 0.3). <u>Interaction niche widths:</u> *σ*_*Pol*_ = 1.7, *σ*_*Her*_ = 2.

At weak trade-offs, varying interaction basal rates (Fig. 5) or niche widths (Fig. S2) mainly affects the composition and stability of the ecological communities, i.e. background colors in Fig. 5 & S2. The selective landscape, characterized by stabilizing vs. runaway selection close to the pollinator vs. herbivore phenotype (as in Fig. 2A), remains essentially unaltered. In particular, increasing the basal rate of pollination eventually selects for phenotypes leading to unstable community dynamics (e.g. Fig. 5a vs. 5b, unbounded growth). Increasing the rate of herbivory restores stability (e.g. Fig. 5b vs. 5d). Note also the narrower set of plant phenotypes allowing coexistence when herbivory becomes more specialized (e.g. Fig. S2a vs. S2c). Close to the herbivore phenotype, the resulting increase in herbivory strength (equation 2) reduces plant density so that pollinators no longer survive. Close to the pollinator phenotype, the resulting decrease in herbivory strength impedes herbivore survival. Plant evolution within the three-species community then provokes the extinction of herbivores. More specialized pollination can restore coexistence, but exposes the community to unstable dynamics (Fig. S2c vs. S2d, unbounded growth).

At strong trade-offs, basal rates or niche widths variations primarily modify the selective landscape – i.e. evolutionary singularities in Fig. 5 & S2 – while the ecological context remains essentially unaffected. Evolutionary dynamics may utterly change. For instance, increasing herbivory can shift the selection regime from stabilizing to disruptive (Fig. 5a vs. 5c), due to the modification of the number of singularities (one vs. three), and of their type (a CSS becoming a branching point). The plant-animal community is significantly modified by the emergence of plant dimorphism, maintained in this case. Modifications of the position of singularities, CSSs in particular, can also have far-reaching implications for the maintained community. A stronger herbivory displaces CSS phenotypes closer to animal extinction thresholds (e.g. Fig. 5a vs. 5c), which implies a fragile coexistence. Given small animal densities, coexistence is indeed vulnerable to perturbations or stochasticity. Stronger pollination, in contrast, displaces the CSS phenotypes away from extinction boundaries (e.g. Fig. 5c vs. 5d).

### Maintenance of plant dimorphism arising from disruptive selection

Disruptive selection on the plant phenotype arises from the interplay of pollination and herbivory. The resulting dimorphism is often temporary, but stable dimorphism is possible in the case of strong trade-offs. Simulating the eco-evolutionary dynamics following the 676 branching points encountered during our second Monte Carlo experiment when constraining the sampling of trade-off intensity, we indeed find that dimorphism is maintained in only 6% of the cases, all occurring at very strong trade-offs (*t*_*h*_ − *t*_*m*_ ≤ 0.5). At such trade-offs, however, the maintenance of dimorphism is relatively frequent, representing around 27% of the cases (details in appendix D). Interestingly, we find that the ratio of pollination to herbivory at branching 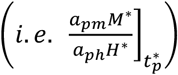 again largely explains the maintenance of dimorphism at these strongest trade-offs (Wilcoxon test, *p*_*value*_ < 10^−10^ & *effect*_*size*_ ≈ 50%, see Fig. 6A). Dimorphism maintenance is associated with stronger herbivory than pollination, which suggests a decisive role of herbivory in the process. We finally show that even a slight amount of similarity-dependent competition suffices to maintain dimorphism in the cases where it is lost (Fig. 6B). To do this, we reformulate our model assumptions to make intraspecific plant competition depend on the phenotypes of competing plant individuals (equation 5). Competition between two plant phenotypes (*t*_*p*1_, *t*_*p*2_) now declines as their niche-overlap (i.e. *t*_*p*1_ − *t*_*p*2_) decreases, *α*_*c*_ and *σ*_*c*_ being the input parameters controlling the proportion and shape of similarity-dependent competition, respectively.

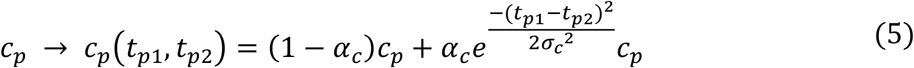

**Fig. 6:**
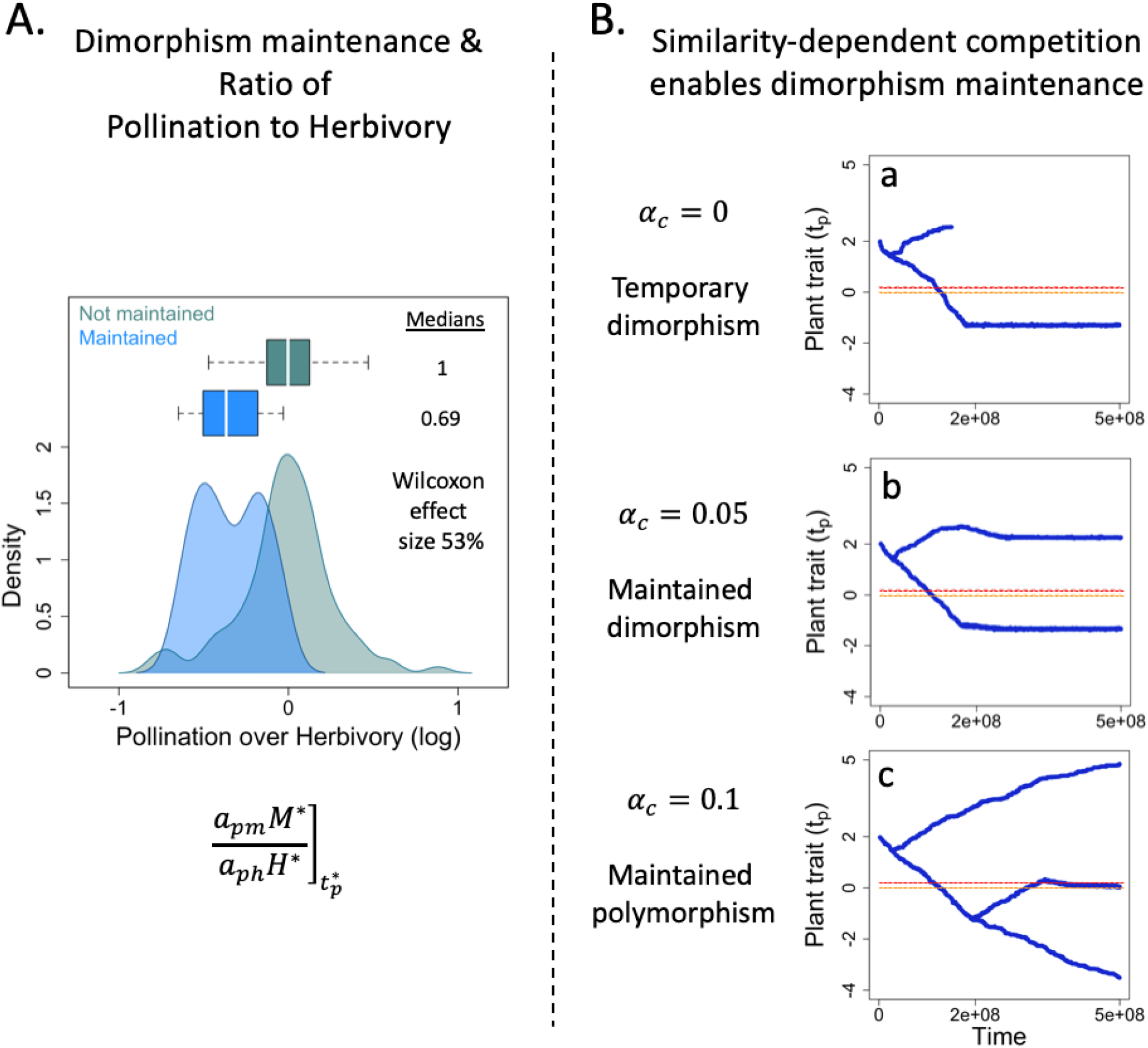
The maintenance of plant dimorphism. A. Ratio of pollination and herbivory at branching explains the maintenance of plant dimorphism. The branchings considered here are those occurring at very strong trade-offs (*t*_*h*_ − *t*_*m*_ ≤ 0.5, 154/676), given that dimorphism maintenance is only possible at such strong trade-offs. Wilcoxon *p*_*value*_ < 10^−10^. **B. Emergence and maintenance of polymorphism are favored by similarity-dependent competition. a. No similarity-dependent competition**. The branching corresponds to that presented in **Fig. 2B.e. b. Five percent of similarity-dependent competition** enables the maintenance of plant dimorphism. **c. Ten percent of similarity-dependent competition** enables secondary branchings, leading to a polymorphic plant population. <u>Ecological parameter set:</u> (*r*_*p*_ = 10, *r*_*m*_ = −1, *r*_*h*_ = −4, *c*_*p*_ = 0.6, *c*_*m*_ = 0.5, *c*_*h*_ = 0.4, *e*_*m*_ = 0.2, *e*_*h*_ = 0.3). <u>Interspecific parameter set:</u> (*a*_*pm*0_ = 5, *a*_*ph*0_ = 7, *σ*_*Pol*_ = 3, *σ*_*Her*_ = 2.8). Animal phenotypes are indicated with orange (pollinator) and red (herbivore) dotted lines: (*t*_*m*_ = 0, *t*_*h*_ = 0.2).

Five percent of similarity-dependence can enable the maintenance of dimorphism (Fig. 6B. a vs. b). Ten percent of similarity-dependence can lead to secondary branching events and the emergence of plant polymorphism (Fig. 6B.c), which suggests a potential role of the interplay between pollination, herbivory and associated niche-overlap competition in the diversification of flowering plant species.

## Discussion

In this article, we investigate how the eco-evolutionary dynamics of plant-pollinator-herbivore communities are driven by the evolution of plant traits under conflicting selection. Conflicting selection arises from the ecological trade-off between attracting pollinators and escaping herbivores which naturally emerges in a trait-matching framework. We find that stronger pollination makes stabilizing selection more prevalent and facilitates coexistence. Stronger herbivory increases the prevalence of runaway selection and threatens coexistence. Importantly, joint selection may be disruptive, leading to plant diversification. Such a diversification requires strong trade-offs, with the strongest trade-offs allowing its long-term maintenance. At weak trade-offs, coexistence is threatened as runaway dynamics are more frequent. Stabilizing selection can still maintain coexistence, but the intrinsic imbalance of plant-animal interactions makes it vulnerable to perturbations. The strength of the ecological trade-off appears as a structuring determinant of community eco-evolutionary dynamics.

We find that pollination fosters the convergence toward the pollinator phenotype – i.e. stabilizing selection – while herbivory favors the divergence from the herbivore phenotype – i.e. runaway selection. Such opposite selection pressures have notably been found acting on flower color (Irwin *et al*. 2003; Frey 2004), shape (Galen & Cuba 2001) or display (Gómez 2005), nectar quantity (Adler & Bronstein 2004) and flowering phenology (Brody 2008; Ehrlén & Münzbergová 2009). Conceptually, all traits that make the discovery of a plant species easier for interacting species – i.e. more “apparent” (Feeny 1976), e.g. high abundance, large size, wide phenology… – may experience ecological trade-offs as they facilitate the discovery by both allies and enemies. A consequence of such trade-offs is that plant phenotypes deviate from optima favored by pollinators (CSSs ≠ *t*_*m*_ in Fig. 5), as illustrated by Ramos & Schiestl (2019). Using experimental evolution, the authors show that plants that evolved in the presence of both pollinators and herbivores were less attractive to pollinators than those that evolved with only pollinators, but still more attractive than plants that evolved with hand-pollination, with or without herbivores. The presence of pollinators thus selected toward plant-pollinator trait matching, but the presence of herbivores reduced such matching by adding a runaway component to selection (equation (3)).

Recent data have highlighted the occurrence of vast insect declines (Hallmann *et al*. 2017; Outhwaite *et al*. 2022). Such declines may be accompanied by changes in the relative frequency of antagonistic vs mutualistic interactions (e.g. herbivores vs pollinators). Given massive pollinator declines (Potts *et al*. 2010) for instance, plant-animal interactions could become increasingly biased towards antagonism. Given our results, this should favor runaway selection thereby leading to plant phenotypes maladapted to pollinators, further accelerating their declines (see also Weinbach et al. 2022). Herbivore extinctions are also expected as a result of more frequent runaway dynamics. Contrary to our assumption of being fixed, animal phenotypes also evolve to match their resource phenotype in natural settings (e.g. Berenbaum and Zangerl 1998; Muchhala 2006). Runaway dynamics are therefore not expected to systematically provoke the extinction of one or both animals, but their ecological outcome will rather depend on the relative speed of evolution of the different species (e.g. evolutionary rescue, Gomulkiewicz and Holt 1995).

More generally, several assumptions that simplify the evolutionary process are made within our modeling framework. The assumption of fixed animal phenotypes might prove to be relevant for some specific communities in which the plant evolves at a much faster rate than the animals owing to higher mutation rates, standing genetic variation, reproduction rates or lifespans. Animal morphologies, preferences, detection abilities or phenologies may however coevolve with plant phenotypes in natural communities (Thompson 2009). Results of this coevolution will depend on the relative speeds of evolution of all interactors, which may lead to many different scenarios whose analyses go beyond the scope of this article. Our work aims at developing an in-depth analytical understanding of the dynamics resulting from the sole plant evolution, before more complex coevolutionary scenarios are studied. We acknowledge that it is not straightforward to extend our results to these scenarios as a higher trait dimensionality can substantially alter evolutionary dynamics (Kisdi 2006; Débarre *et al*. 2014). Another assumption of our framework is that plant evolution has no other costs than those ensuing from pollination and herbivory. We used this assumption to better highlight the direct implications of ecological trade-offs. This might be a reasonable approximation for specific traits whose selective pressures essentially arise from these plant-animal interactions (e.g. presumably flower color in *Raphanus sativus*, Irwin et al. 2003), but plant traits are highly multifunctional in general. They are indeed often also involved in other types of interactions (Strauss & Whittall 2006) or processes such as stress tolerance (Sack & Buckley 2020). A direct consequence is that phenotypes are usually constrained within an ecologically viable space, which notably sets a limit to runaway dynamics, mitigating the likelihood of extinctions.

In line with the empirical evidence so far (Strauss *et al*. 2002; Strauss & Irwin 2004), our study identifies the strength of ecological trade-off as a key driver of plant-animal eco-evolutionary dynamics. We underline that trade-off intensity affects the maintenance of coexistence, as well as the nature of joint selection, in particular the potential for disruptive selection and plant diversification.

We find that coexistence is fragile or destroyed at weak trade-offs. Runaway dynamics are more frequent (Fig. 4A). Evolutionary-induced unstable dynamics become possible (Fig. 5b&d). In the vicinity of the pollinator phenotype, the plant phenotype is more likely to stabilize after the extinction of herbivores, as indicated by the CSSs leaving the coexistence area (Fig. 2A). When stabilizing selection occurs before herbivore extinction, their low density makes them vulnerable to demographic stochasticity, Allee effects or external disturbances. Because weak ecological trade-offs are associated with fragile plant-pollinator-herbivore coexistence, they should be seldom encountered in natural communities. Accordingly, shared animal preferences for plant phenotype are most often reported in the empirical literature (Adler & Bronstein 2004; Strauss & Whittall 2006).

Selection can turn disruptive under stronger ecological trade-offs. We stress that disruptive selection here arises from the joint selection of pollination and herbivory. In the absence of either one interaction, branching is not possible here, as evolutionary dynamics would run away from the herbivore phenotype in the absence of pollination, and would stabilize at the pollinator phenotype in the absence of herbivory. Diversification we uncover here is thus fundamentally different from the one uncovered in previous trait-matching models that consider the two interactions separately (e.g. Yoder and Nuismer 2010; Maliet et al. 2020). While dimorphism can only be maintained when the trade-off is sufficiently strong, long-term maintenance is frequent in such instances. Polymorphism then yields two contrasted phenotypes: one that has many allies and enemies, while the other one has few interactions. Interestingly, such a pattern has been frequently reported in the empirical literature. In *Primula farinosa*, the fitness advantage of long-scaped phenotypes resulting from higher pollinator visitation rates is balanced by higher levels of fruit predation, allowing the maintenance of dimorphism in scape length (Ehrlén *et al*. 2002). The maintenance of color dimorphism in *Raphanus sativus* (Irwin *et al*. 2003) and *Claytonia virginica* (Frey 2004) is similarly supported by animal species preferentially interacting with the same plant phenotype. Field studies of tripartite networks also report a positive correlation between the number (Sauve *et al*. 2016b) or strength (Melián *et al*. 2009) of plant-pollinator and plant-herbivore interactions of plant species, indicating a rather widespread pattern. This mechanism of diversification under balanced interaction strengths may also apply on longer evolutionary timescales. Using a phylogenetic analysis on 32 *Nicotiana* species, Adler *et al*. (2012) show that variations in nicotine defenses among species are largely negatively correlated to pollinator reliance. These tobacco species are therefore either well-defended with few pollinators, or lowly defended and relying on pollination.

We find that disruptive selection requires a local balance between pollination and herbivory (Fig 3), while its prevalence within the selective landscape increases with the strength of herbivory (Fig. 4B). This is not contradictory as those aspects describe selection at two different scales. To put it simply, disruptive selection can be expected in any three-species community when plant-animal interactions are well-balanced, but communities in which herbivores are more ravenous (*a*_*ph*0_ ↗) will more likely exhibit such evolutionary dynamics. Dimorphism is also more likely to be permanent with stronger herbivory (Fig. 6A), further emphasizing its decisive role in diversification processes. Previous theoretical works that contrast interaction types but one at a time have also highlighted the crucial role of antagonisms for diversification (Kopp & Gavrilets 2006; Yoder & Nuismer 2010; Maliet *et al*. 2020). Given these results, we note that observed declines in the diversity and abundances of herbivore species (Sánchez-Bayo & Wyckhuys 2019; Atwood *et al*. 2020) may have far-reaching consequences if they lead to future reductions in diversification rates.

Overall, dimorphism is often temporary within our framework. While we assume constant competition rates in order to focus on how trophic vs mutualistic interactions shape eco-evolutionary dynamics, similarity-dependent competition is likely to occur in natural communities (Macarthur & Levins 1967). Here, similar plants would share pollinators, which could lead to strong pollination niche-overlap competition for instance due to the dilution of pollen between plant species (Morales & Traveset 2008; Mitchell *et al*. 2009). Similarly, shared herbivores would increase apparent competition. Plant traits can also be simultaneously involved in both competitive and plant-animal interactions, either directly (e.g. phenological traits, Schwinning & Kelly 2013) or due to genetic correlations (e.g. plant size, Carmona *et al*. 2011). Accounting for a small amount of niche-overlap competition suffice to maintain dimorphism in our model, and enhances the potential for further diversification (Fig. 6B). Reproductive isolation may then evolve as the morph that interacts weakly with animal species acquires new pollinators species (Baack *et al*. 2015), or evolves self-fertilization (Bodbyl Roels & Kelly 2011). Disruptive selection from the interplay of pollination, herbivory and niche-overlap competition can therefore be construed as a first step toward modeling the evolutionary emergence of plant-pollinator-herbivore networks, which would allow new insights into the eco-evolutionary processes supporting the diversity of natural communities.

## Supporting information

Supplementary details, notably including the derivation of mathematical expressions and demonstrations, are provided online (directly on BioRxiv, or on Zenodo via https://doi.org/10.5281/zenodo.11263481). It consists of 2 additional figures (S1, S2), one table (S1) and 4 appendices (A, B, C & D).

## Data availability statement

The codes used to conduct the Monte Carlo numerical experiments, as well as that used to simulate the community eco-evolutionary dynamics, are available in Zenodo (https://doi.org/10.5281/zenodo.11263481).

## Acknowledgements

The authors would like to thank Dr. Sebastien Lion for his helpful feedback on the manuscript.

## References

Adler, L.S. & Bronstein, J.L. (2004). Attracting antagonists: Does floral nectar increase leaf herbivory? Ecology, 85, 1519–1526.

Adler, L.S., Seifert, M.G., Wink, M. & Morse, G.E. (2012). Reliance on pollinators predicts defensive chemistry across tobacco species. Ecol. Lett., 15, 1140–1148.

Armbruster, W.S. (1997). Exaptations Link Evolution of Plant-Herbivore and Plant-Pollinator Interactions: A Phylogenetic Inquiry. Ecology, 78, 1661.

Atwood, T.B., Valentine, S.A., Hammill, E., McCauley, D.J., Madin, E.M.P., Beard, K.H., et al. (2020). Herbivores at the highest risk of extinction among mammals, birds, and reptiles. Sci. Adv., 6, eabb8458.

Baack, E., Melo, M.C., Rieseberg, L.H. & Ortiz-Barrientos, D. (2015). The origins of reproductive isolation in plants. New Phytol., 207, 968–984.

Becerra, J.X., Noge, K. & Venable, D.L. (2009). Macroevolutionary chemical escalation in an ancient plant-herbivore arms race. Proc. Natl. Acad. Sci. U. S. A., 106, 18062–18066.

Berenbaum, M.R. & Zangerl, A.R. (1998). Chemical phenotype matching between a plant and its insect herbivore. Proc. Natl. Acad. Sci. U. S. A., 95, 13743–13748.

Bodbyl Roels S.A. & Kelly, J.K. (2011). Rapid evolution caused by pollinator loss in Mimulus guttatus. Evolution (N. Y)., 65, 2541–2552.

Brody, A.K. (2008). Effects of Pollinators, Herbivores, and Seed Predators on Flowering Phenology. Ecology, 78, 1624–1631.

Carmona, D., Lajeunesse, M.J. & Johnson, M.T.J. (2011). Plant traits that predict resistance to herbivores. Funct. Ecol., 25, 358–367.

Chamberlain, S.A., Bronstein, J.L. & Rudgers, J.A. (2014). How context dependent are species interactions? Ecol. Lett., 17, 881–890.

Craine, J.M. & Dybzinski, R. (2013). Mechanisms of plant competition for nutrients, water and light. Funct. Ecol., 27, 833–840.

Dawkins, R. & Krebs, J.R. (1979). Arms races between and within species. Proc. R. Soc. B Biol. Sci., 205, 489– 511.

Débarre, F., Nuismer, S.L. & Doebeli, M. (2014). Multidimensional (Co)evolutionary stability. Am. Nat., 184, 158–171.

Dieckmann, U. & Law, R. (1996). The dynamical theory of coevolution: a derivation from stochastic ecological processes. J. Math. Biol., 34, 579–612.

Ehrlén, J., Käck, S. & Ågren, J. (2002). Pollen limitation, seed predation and scape length in Primula farinosa. Oikos, 97, 45–51.

Ehrlén, J. & Münzbergová, Z. (2009). Timing of flowering: Opposed selection on different fitness components and trait covariation. Am. Nat., 173, 819–830.

Ehrlich, P.R. & Raven, P.H. (1964). Butterflies and Plants: A Study in Coevolution. Evolution (N. Y)., 18, 586.

Eshel, I. (1983). Evolutionary and continuous stability. J. Theor. Biol., 103, 99–111.

Farrell, B.D., Dussourd, D.E. & Mitter, C. (1991). Escalation of plant defense: do latex and resin canals spur plant diversification? Am. Nat., 138, 881–900.

Feeny, P. (1976). Plant apparency and chemical defense. In: Biochemical interaction between plants and insects. Springer, Boston, MA, pp. 1–40.

Fontaine, C., Guimarães Jr, P.R., Kéfi, S., Loeuille, N., Memmott, J., van der Putten, W.H., et al. (2011). The ecological and evolutionary implications of merging different types of networks. Ecol. Lett., 14, 1170–1181.

Frey, F.M. (2004). Opposing natural selection from herbivores and pathogens may maintain floral-color variation in Claytonia virginica (Portulacaceae). Evolution (N. Y)., 58, 2426–2437.

Galen, C. & Cuba, J. (2001). Down the tube: Pollinators, predators, and the evolution of flower shape in the alpine skypilot, Polemonium viscosum. Evolution (N. Y)., 55, 1963–1971.

Geritz, S.A.H., Metz, J.A.J., Kisdi É. & Meszéna, G. (1997). Dynamics of adaptation and evolutionary branching. Phys. Rev. Lett., 78, 2024–2027.

Gómez, J.M. (2005). Non-additive effects of herbivores and pollinators on Erysimum mediohispanicum (Cruciferae) fitness. Oecologia, 143, 412–418.

Gómez, J.M., Iriondo, J.M. & Torres, P. (2023). Modeling the continua in the outcomes of biotic interactions. Ecology, 104, e3995.

Gomulkiewicz, R. & Holt, R.D. (1995). When does Evolution by Natural Selection Prevent Extinction? Evolution (N. Y)., 49, 201–207.

Grant, V. (1949). Pollination Systems as Isolating Mechanisms in Angiosperms. Evolution (N. Y)., 3, 82–97.

Griese, E., Caarls, L., Bassetti, N., Mohammadin, S., Verbaarschot, P., Bukovinszkine’Kiss, G., et al. (2021). Insect egg-killing: a new front on the evolutionary arms-race between brassicaceous plants and pierid butterflies. New Phytol., 230, 341–353.

Hahn, M. & Brühl, C.A. (2016). The secret pollinators: an overview of moth pollination with a focus on Europe and North America. Arthropod. Plant. Interact., 10, 21–28.

Hallmann, C.A., Sorg, M., Jongejans, E., Siepel, H., Hofland, N., Schwan, H., et al. (2017). More than 75 percent decline over 27 years in total flying insect biomass in protected areas. PLoS One, 12.

Hernández-Hernández, T. & Wiens, J.J. (2020). Why are there so many flowering plants? A multiscale analysis of plant diversification. Am. Nat., 195, 948–963.

Hodges, S.A. & Arnold, M.L. (1995). Spurring plant diversification: Are floral nectar spurs a key innovation? Proc. R. Soc. B Biol. Sci., 262, 343–348.

Holt, R.D. (1977). Predation, apparent competition, and the structure of prey communities. Theor. Popul. Biol., 12, 197–229.

Hutson, V. & Law, R. (1985). Permanent coexistence in general models of three interacting species. J. Math. Biol., 21, 285–298.

Irwin, R.E., Strauss, S.Y., Storz, S., Emerson, A. & Guibert, G. (2003). The Role of Herbivores in the Maintenance of a Flower Color Polymorphism in Wild Radish. Ecology, 84, 1733–1743.

De Jager, M.L. & Peakall, R. (2019). Experimental examination of pollinator-mediated selection in a sexually deceptive orchid. Ann. Bot., 123, 347–354.

Kessler, A., Halitschke, R. & Poveda, K. (2011). Herbivory-mediated pollinator limitation: Negative impacts of induced volatiles on plant-pollinator interactions. Ecology, 92, 1769–1780.

Kiester, A.R., Lande, R. & Schemske, D.W. (1984). Models of coevolution and speciation in plants and their pollinators. Am. Nat., 124, 220–243.

Kisdi É. (2006). Trade-off geometries and the adaptive dynamics of two co-evolving species. Evol. Ecol. Res., 8, 959–973.

Kopp, M. & Gavrilets, S. (2006). Multilocus genetics and the coevolution of quantitative traits. Evolution (N. Y)., 60, 1321–1336.

Macarthur, R. & Levins, R. (1967). The Limiting Similarity, Convergence, and Divergence of Coexisting Species. Am. Nat., 101, 377–385.

Maliet, O., Loeuille, N. & Morlon, H. (2020). An individual-based model for the eco-evolutionary emergence of bipartite interaction networks. Ecol. Lett., 23, 1623–1634.

Mauricio, R. & Rausher, M.D. (1997). Experimental manipulation of putative selective agents provides evidence for the role of natural enemies in the evolution of plant defense. Evolution (N. Y)., 51, 1435–1444.

Maynard Smith, J. & Price, G.R. (1973). The logic of animal conflict. Nature, 246, 15–18.

Melián, C.J., Bascompte, J., Jordano, P. & Křivan, V. (2009). Diversity in a complex ecological network with two interaction types. Oikos, 118, 122–130.

Metz, J.A.J., Nisbet, R.M. & Geritz, S.A.H. (1992). How should we define “fitness” for general ecological scenarios? Trends Ecol. Evol., 7, 198–202.

Mitchell, R.J., Flanagan, R.J., Brown, B.J., Waser, N.M. & Karron, J.D. (2009). New frontiers in competition for pollination. Ann. Bot., 103, 1403–1413.

Morales, C.L. & Traveset, A. (2008). Interspecific pollen transfer: Magnitude, prevalence and consequences for plant fitness. CRC. Crit. Rev. Plant Sci., 27, 221–238.

Mougi, A. & Kondoh, M. (2014). Instability of a hybrid module of antagonistic and mutualistic interactions. Popul. Ecol., 56, 257–263.

Muchhala, N. (2006). Nectar bat stows huge tongue in its rib cage. Nature, 444, 701–702.

Ollerton, J., Winfree, R. & Tarrant, S. (2011). How many flowering plants are pollinated by animals? Oikos, 120, 321–326.

Outhwaite, C.L., McCann, P. & Newbold, T. (2022). Agriculture and climate change are reshaping insect biodiversity worldwide. Nature, 605, 97–102.

Parachnowitsch, A.L. & Kessler, A. (2010). Pollinators exert natural selection on flower size and floral display in Penstemon digitalis. New Phytol., 188, 393–402.

Potts, S.G., Biesmeijer, J.C., Kremen, C., Neumann, P., Schweiger, O. & Kunin, W.E. (2010). Global pollinator declines: Trends, impacts and drivers. Trends Ecol. Evol., 25, 345–353.

Ramos, S.E. & Schiestl, F.P. (2019). Rapid plant evolution driven by the interaction of pollination and herbivory. Science, 364, 193–196.

Sack, L. & Buckley, T.N. (2020). Trait multi-functionality in plant stress response. Integr. Comp. Biol., 60, 98– 112.

Sahli, H.F. & Conner, J.K. (2011). Testing for conflicting and nonadditive selection: Floral adaptation to multiple pollinators through male and female fitness. Evolution (N. Y)., 65, 1457–1473.

Sánchez-Bayo, F. & Wyckhuys, K.A.G. (2019). Worldwide decline of the entomofauna: A review of its drivers. Biol. Conserv., 232, 8–27.

Sargent, R.D. (2004). Floral symmetry affects speciation rates in angiosperms. Proc. R. Soc. B Biol. Sci., 271, 603–608.

Sauve, A.M.C., Fontaine, C. & Thébault, E. (2014). Structure-stability relationships in networks combining mutualistic and antagonistic interactions. Oikos, 123, 378–384.

Sauve, A.M.C., Fontaine, C. & Thébault, E. (2016a). Stability of a diamond-shaped module with multiple interaction types. Theor. Ecol., 9, 27–37.

Sauve, A.M.C., Thébault, E., Pocock M.J.O. & Fontaine, C. (2016b). How plants connect pollination and herbivory networks and their contribution to community stability. Ecology, 97, 908–917.

Schwinning, S. & Kelly, C.K. (2013). Plant competition, temporal niches and implications for productivity and adaptability to climate change in water-limited environments. Funct. Ecol., 27, 886–897.

Strauss, S.Y. & Irwin, R.E. (2004). Ecological and Evolutionary Consequences of Multispecies Plant-Animal Interactions. Annu. Rev. Ecol. Evol. Syst., 35, 435–466.

Strauss, S.Y., Irwin, R.E. & Lambrix, V.M. (2004). Optimal defence theory and flower petal colour predict variation in the secondary chemistry of wild radish. J. Ecol., 92, 132–141.

Strauss, S.Y., Rudgers, J.A., Lau, J.A. & Irwin, R.E. (2002). Direct and ecological costs of resistance to herbivory. Trends Ecol. Evol., 17, 278–285.

Strauss, S.Y. & Whittall, J.B. (2006). Non-pollinator agents of selection on floral traits. In: Ecology and evolution of flowers (eds. Harder L.D. & Barrett S.C.H.). Oxford University Press on Demand, New York, NY, USA, pp. 120–138.

Thébault, E. & Fontaine, C. (2010). Stability of ecological communities and the architecture of mutualistic and trophic networks. Science, 329, 853–856.

Theis, N., Barber, N.A., Gillespie, S.D., Hazzard, R. V. & Adler, L.S. (2014). Attracting mutualists and antagonists: Plant trait variation explains the distribution of specialist floral herbivores and pollinators on crops and wild gourds. Am. J. Bot., 101, 1314–1322.

Thompson, J.N. (1988). Variation in interspecific interactions. Annu. Rev. Ecol. Syst. Vol. 19, 19, 65–87.

Thompson, J.N. (2009). The coevolving web of life. Am. Nat., 173, 125–140.

Weinbach, A., Loeuille, N. & Rohr, R.P. (2022). Eco-evolutionary dynamics further weakens mutualistic interaction and coexistence under population decline. Evol. Ecol., 36, 373–387.

Whittall, J.B. & Hodges, S.A. (2007). Pollinator shifts drive increasingly long nectar spurs in columbine flowers. Nature, 447, 706–709.

Yacine, Y. & Loeuille, N. (2022). Stable coexistence in plant-pollinator-herbivore communities requires balanced mutualistic vs antagonistic interactions. Ecol. Modell., 465, 109857.

Yoder, J.B. & Nuismer, S.L. (2010). When Does Coevolution Promote Diversification? Am. Nat., 176, 802–817.

