## Supplementary Information for "Attracting pollinators vs escaping herbivores: eco-evolutionary dynamics of plants confronted with an ecological trade-off"

### Supporting Material

**Figure S1**……………………………………………………...2

**Table S1**………………………………..………………..……..3

**Figure S2**……………………………………………………..……..4

**Appendix A:** Analytical investigation of plant-pollinator-herbivore eco-evolutionary dynamics……………………………………...5

I. Ecological equilibria………………………………………...…….5

II. Adaptive dynamics ……………………………………..……….6

**Appendix B:** Setting and exploring the parameter space…..................10

I. Choosing parameter values……………………………..…………...10

II. Monte Carlo experiments……………………..………..……………12

**Appendix C:** Robustness to the variation of the ecological parameter set…………………………………………………………16

I. Relationship between type of selection and relative strength of pollination vs. herbivory …………………………………………………….....16

II. Effect of interspecific parameters on the selective landscape...…………………..........................................................................17

**Appendix D:** Joint selection and the emergence of plant dimorphism……………………………………………………..21

Eco-evolutionary landscape associated with Fig. 2B.f

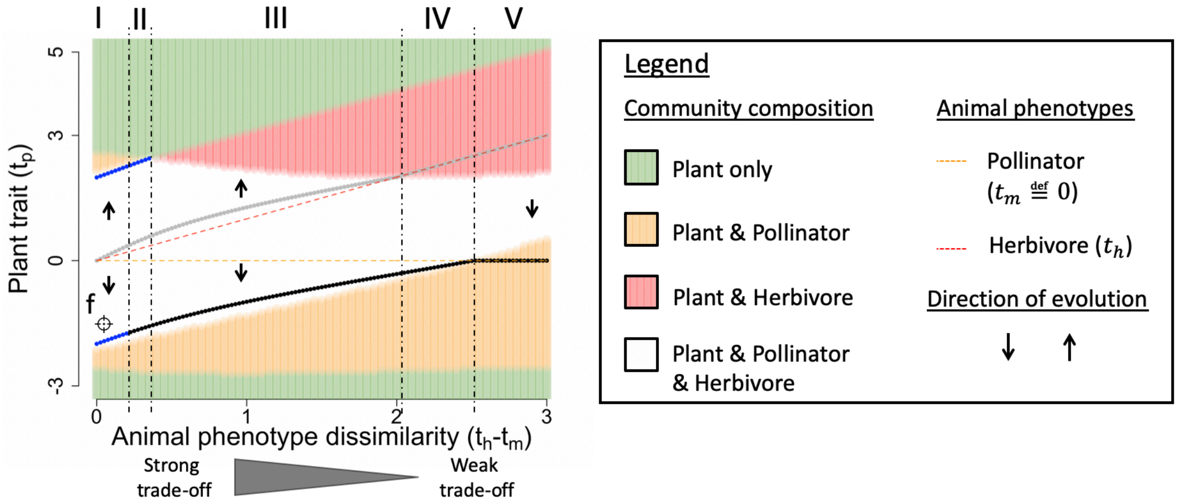


**Fig. S1: Eco-evolutionary landscape associated with the dynamics of maintained dimorphism figured in Fig. 2B.f.** In region I, selection is disruptive. In region II, selection is still disruptive when the plan phenotype is above the repellor, but stabilizing below the repellor (i.e. two basins of attraction separated by the repellor singularity). In region III, selection remains stabilizing for phenotypes below the repellor, but turns runaway above the repellor. Subsequent runaway dynamics lead to the extinction of pollinators, then of herbivores. In region IV, the basin of attraction associated with stabilizing dynamics encompasses all the coexistence area (white). Finally, selection is runaway in region V. Eco-evolutionary dynamics then lead to the extinction of herbivores, leading to a perfectly matching plant-pollinator community. Ecological parameter set: ($r_p = 10, r_m = -1, r_h = -4, c_p = 0.6, c_m = 0.5, c_h = 0.4, e_m = 0.2, e_h = 0.3$). Interspecific parameter set ( $a_{pm0} = 5, a_{ph0} = 9, \sigma_{Pol} = 1.7, \sigma_{Her} = 2$)

| Ratio of pollination to herbivory | Expression | Median per type of selection | | | Kruskal-Wallis effect size |
|---|---|---|---|---|---|
| | | Runaway | Disruptive | Stabilizing | |
| Interaction strengths and animal densities | $\left.\frac{a_{pm}M^*}{a_{ph}H^*}\right]_{t_p^*}$ | 0.13 | 0.92 | 3.26 | 0.674 |
| Interaction strengths | $\left.\frac{a_{pm}}{a_{ph}}\right]_{t_p^*}$ | 0.4 | 0.49 | 1.36 | 0.632 |
| Animal densities | $\left.\frac{M^*}{H^*}\right]_{t_p^*}$ | 0.32 | 1.74 | 2.42 | 0.565 |

**Table S1: Results of the statistical analysis for the three ratios of pollination to herbivory tested.** The ratios are calculated at the evolutionary singularities $t_p^*$. Sample size (interspecific parameter sets): 10000, which resulted in 7814 Stabilizing, 2762 Runaway and 1122 Disruptive. All statistical tests were highly significant, with a $p_{value}$ below $2.2\ 10^{-16}$. Ecological parameter set: ($r_p = 10, r_m = -1, r_h = -4, c_p = 0.6, c_m = 0.5, c_h = 0.4, e_m = 0.2, e_h = 0.3$).

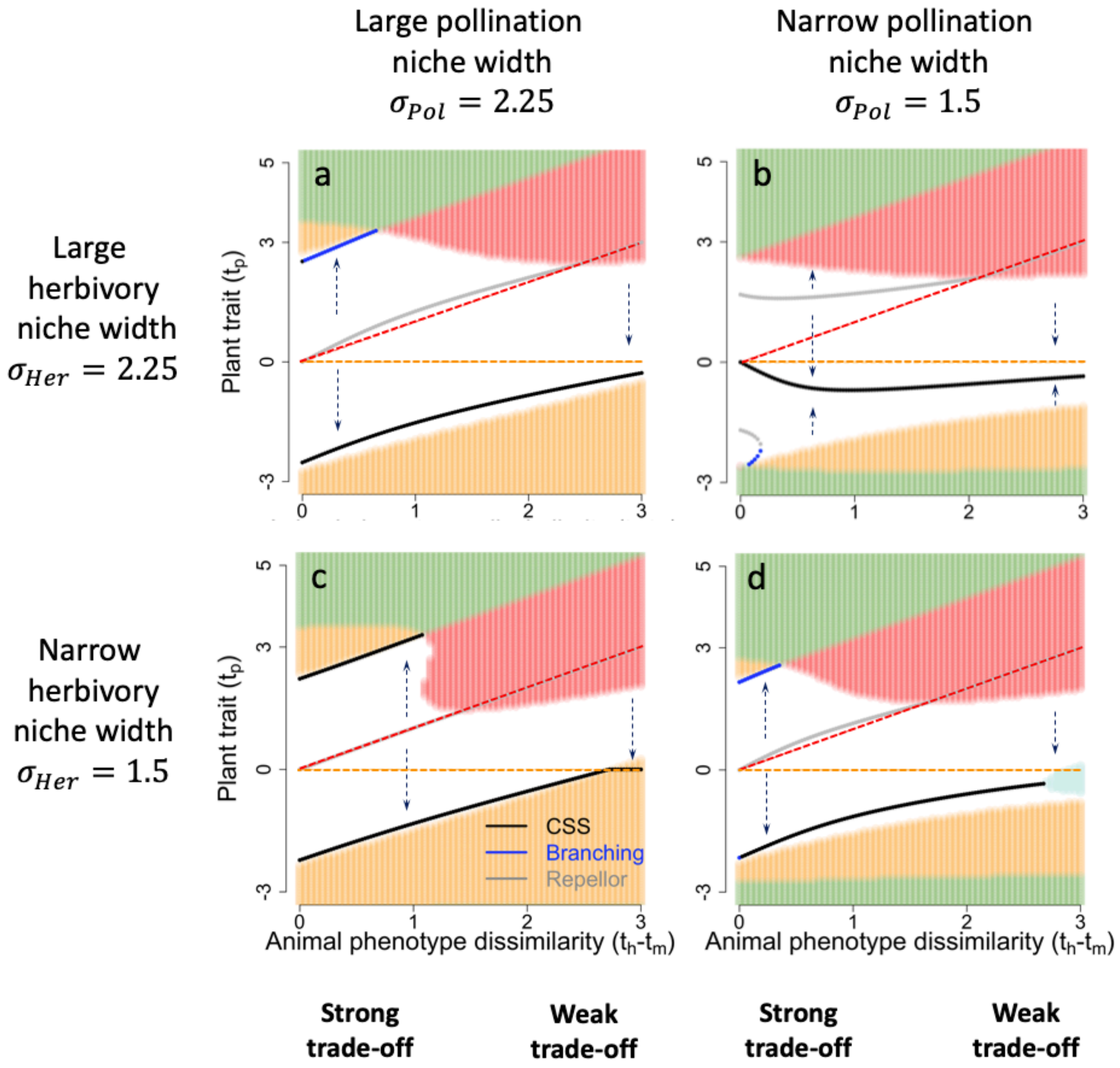


**Fig. S2: Effect of the niche widths of plant-animal interactions on community eco-evolutionary dynamics.** At strong trade-offs, eco-evolutionary dynamics are primarily altered by changes affecting evolutionary singularities. At weak trade-offs, the alteration is mainly mediated by changes in the composition of the ecological community (i.e. background colors). Orange and red dotted lines indicate the pollinator and herbivore phenotype. Arrows indicate evolutionary trajectories and background colors indicate community composition as in **Fig. 5.** Ecological parameter set: $(r_p = 10, r_m = -1, r_h = -4, c_p = 0.6, c_m = 0.5, c_h = 0.4, e_m = 0.2, e_h = 0.3)$. Basal rates for plant-animal interactions: $a_{pm0} = 5,\ a_{ph0} = 9$.

### Appendix A: Analytical investigation of plant-pollinator-herbivore eco-evolutionary dynamics

#### I. Ecological equilibria

Population dynamics reach an ecological equilibrium when the three population growth rates vanish.

$$\begin{cases} \dfrac{dP}{dt} \stackrel{\text{def}}{=} P\left(r_p - c_p P + a_{pm}M - a_{ph}H\right) = 0 \\ \dfrac{dM}{dt} \stackrel{\text{def}}{=} M(r_m - c_m M + e_m a_{pm} P) = 0 \\ \dfrac{dH}{dt} \stackrel{\text{def}}{=} H\left(r_h - c_h H + e_h a_{ph} P\right) = 0 \end{cases}$$

There are eight possible equilibria as each species can either be present or absent at ecological equilibrium. The goal of the present paper is to study the evolutionary dynamics of the plant phenotype within a plant-pollinator-herbivore community. We are thus mostly interested in the coexistence equilibrium, i.e. the ecological equilibrium characterized by the presence of all three species at densities $P^*, M^*, H^*$.

$$\begin{cases} P^* = \dfrac{c_h c_m r_p + c_h a_{pm} r_m - c_m a_{ph} r_h}{c_h c_m c_p - c_h e_m a_{pm}{}^2 + c_m e_h a_{ph}{}^2} \\ M^* = \dfrac{c_h e_m a_{pm} r_p + (c_p c_h + e_h a_{ph}{}^2) r_m - e_m a_{pm} a_{ph} r_h}{c_h c_m c_p - c_h e_m a_{pm}{}^2 + c_m e_h a_{ph}{}^2} \\ H^* = \dfrac{c_m e_h a_{ph} r_p + e_h a_{pm} a_{ph} r_m + (c_p c_m - e_m a_{pm}{}^2) r_h}{c_h c_m c_p - c_h e_m a_{pm}{}^2 + c_m e_h a_{ph}{}^2} \end{cases}$$

Moreover, assuming coexistence is feasible (i.e. $P^*, M^*, H^* > 0$), its stability depends on the satisfaction of two inequalities (Ruth-Hurwitz):

$$\left(c_h c_m c_p - c_h e_m a_{pm}{}^2 + c_m e_h a_{ph}{}^2\right) > 0$$

$$\left(c_p P^* + c_m M^* + c_h H^*\right)\left(P^* M^* \left(c_p c_m - e_m a_{pm}^2\right) + P^* H^* \left(c_p c_h + e_h a_{ph}^2\right) + M^* H^* c_m c_h\right) - P^* M^* H^* \left(c_h c_m c_p - c_h e_m a_{pm}{}^2 + c_m e_h a_{ph}{}^2\right) > 0$$

The analytical expressions for the other equilibria, as well as the conditions of their stability, can be found elsewhere (Yacine and Loeuille 2022).

### II. Adaptive dynamics

#### II.1 Adaptive dynamics framework

Within a monomorphic plant population with phenotype ($t_p$), we consider the appearance of a mutant with phenotype ($t_p'$). Assuming the mutant is initially rare so that the environment (i.e. $P^*, M^*, H^*$) is set by the resident phenotype ($t_p$), the long-term per capita growth rate of the mutant ($P'$) corresponds to its invasion fitness $w(t_p', t_p)$.

$$w(t_p', t_p) \stackrel{\text{def}}{=} \left.\frac{dP'}{P'dt}\right]_{P' \ll P^*} = r_p - c_p P^* + a_{pm}(t_p')M^* - a_{ph}(t_p')H^*$$

$$\Leftrightarrow w(t_p', t_p) = \left(a_{pm}(t_p') - a_{pm}(t_p)\right) M^* - \left(a_{ph}(t_p') - a_{ph}(t_p)\right) H^* \qquad (A.1)$$

The last equality results from the ecological equilibrium. A positive fitness of invasion indicates that the mutant invades and replaces the resident plant population, while it goes extinct otherwise. Assuming mutations of small phenotypic amplitude, the rate of this trait substitution sequence is proportional to the selection gradient (Dieckmann and Law 1996).

$$\textit{Selection gradient} \stackrel{\text{def}}{=} \left.\frac{\partial w(t_p', t_p)}{\partial t_p'}\right]_{t_p' = t_p} = -(t_p - t_m)\frac{a_{pm}(t_p)M^*}{\sigma_{Pol}^2} + (t_p - t_h)\frac{a_{ph}(t_p)H^*}{\sigma_{Her}^2} \qquad (A.2)$$

Evolutionary singularities ($t_p^*$) correspond to phenotypes that nullify the selection gradient.

$$0 = -(t_p^* - t_m)\frac{a_{pm}(t_p^*)M^*}{\sigma_{Pol}^2} + (t_p^* - t_h)\frac{a_{ph}(t_p^*)H^*}{\sigma_{Her}^2}$$

$$\Leftrightarrow \frac{t_p^* - t_h}{t_p^* - t_m} = \frac{a_{pm}(t_p^*)M^* / \sigma_{Pol}^2}{a_{ph}(t_p^*)H^* / \sigma_{Her}^2} \qquad (A.3)$$

As indicated in the main text, the position and nature of evolutionary singularities enable the full characterization of evolutionary dynamics (as long as the plant population remains monomorphic). Note that no singularity can occur in the interval $[t_m, t_p]$ as this would imply a negative ratio $\frac{t_p^* - t_h}{t_p^* - t_m}$, which would never possibly satisfy equation (A.3) whose right side is positive. Singularities are classified into Continuously Stable Strategies (CSS), Branching points (BP) or Repellors according to two properties: invasibility (equation A.4) and convergence (equation A.5). Invasibility specifies whether the singularity may be invaded by nearby mutants (non-invasible: CSS; invasible: BP, Repellor). Convergence indicates that the trait evolves toward the singularity in its vicinity (convergent: CSS, BP; non-convergent: Repellor).

| | |
|---|---|
| $$\left.\frac{\partial^2 w(t_p', t_p)}{\partial {t_p}'^2}\right]_{t_p'=t_p=t_p^*} \stackrel{\text{def}}{=} -M^* \frac{a_{pm}(t_p^*)}{\sigma_{Pol}^2}\left[1 - \frac{(t_p^* - t_m)^2}{\sigma_{Pol}^2}\right] + H^* \frac{a_{ph}(t_p^*)}{\sigma_{Her}^2}\left[1 - \frac{(t_p^* - t_h)^2}{\sigma_{Her}^2}\right] > 0$$ | $(A.4)$ |

| | |
|---|---|
| $$\left.\frac{\left.\frac{\partial w(t_p', t_p)}{\partial {t_p}'}\right]_{t_p'=t_p}}{dt_p}\right]_{t_p=t_p^*} \stackrel{\text{def}}{=} -(t_p^* - t_m)\frac{a_{pm}(t_p^*)}{\sigma_{Pol}^2}\left.\frac{dM^*}{dtp}\right]_{t_p=t_p^*} + (t_p^* - t_h)\frac{a_{ph}(t_p^*)}{\sigma_{Her}^2}\left.\frac{dH^*}{dtp}\right]_{t_p=t_p^*} - M^* \frac{a_{pm}(t_p^*)}{\sigma_{Pol}^2}\left[1 - \frac{(t_p^* - t_m)^2}{\sigma_{Pol}^2}\right] + H^* \frac{a_{ph}(t_p^*)}{\sigma_{Her}^2}\left[1 - \frac{(t_p^* - t_h)^2}{\sigma_{Her}^2}\right] < 0$$ | $(A.5)$ |

#### II.2 Analytical investigation of the condition for invasibility

Here, we demonstrate inequality (4) of the main document.
Using equation (A.3) (i.e. the fact that evolutionary singularities nullify the selection gradient):

$$\frac{t_p^* - t_h}{t_p^* - t_m} = \frac{a_{pm}(t_p^*)M^*\sigma_{Her}^2}{a_{ph}(t_p^*)H^*\sigma_{Pol}^2}$$

We have:

$$t_p^* = t_m + \frac{\frac{a_{ph}(t_p^*)H^*}{\sigma_{Her}^2}(t_h - t_m)}{\frac{a_{ph}(t_p^*)H^*}{\sigma_{Her}^2} - \frac{a_{pm}(t_p^*)M^*}{\sigma_{Pol}^2}}$$

Or alternatively

$$t_p^* = t_h + \frac{\frac{a_{pm}(t_p^*)M^*}{\sigma_{Pol}^2}(t_h - t_m)}{\frac{a_{ph}(t_p^*)H^*}{\sigma_{Her}^2} - \frac{a_{pm}(t_p^*)M^*}{\sigma_{Pol}^2}}$$

Replacing $t_p^*$ by one of these two expressions in the condition for invasibility (equation A.4) gives:

$$Invasibility \Leftrightarrow -M^* \frac{a_{pm}(t_p^*)}{\sigma_{Pol}^2}\left[1 - \frac{1}{\sigma_{Pol}^2}\left(\frac{\frac{a_{ph}(t_p^*)H^*}{\sigma_{Her}^2}(t_h - t_m)}{\frac{a_{ph}(t_p^*)H^*}{\sigma_{Her}^2} - \frac{a_{pm}(t_p^*)M^*}{\sigma_{Pol}^2}}\right)^2\right]$$
$$+ H^* \frac{a_{ph}(t_p^*)}{\sigma_{Her}^2}\left[1 - \frac{1}{\sigma_{Her}^2}\left(\frac{\frac{a_{pm}(t_p^*)M^*}{\sigma_{Pol}^2}(t_h - t_m)}{\frac{a_{ph}(t_p^*)H^*}{\sigma_{Her}^2} - \frac{a_{pm}(t_p^*)M^*}{\sigma_{Pol}^2}}\right)^2\right] > 0$$

$$\Leftrightarrow -\frac{a_{pm}(t_p^*)M^*}{\sigma_{Pol}^2} + \frac{a_{ph}(t_p^*)H^*}{\sigma_{Her}^2}$$
$$+ \frac{[a_{pm}(t_p^*)M^*][a_{ph}(t_p^*)H^*](t_h - t_m)^2}{\sigma_{Pol}^4\sigma_{Her}^4\left(\frac{a_{ph}(t_p^*)H^*}{\sigma_{Her}^2} - \frac{a_{pm}(t_p^*)M^*}{\sigma_{Pol}^2}\right)^2}[a_{ph}(t_p^*)H^* - a_{pm}(t_p^*)M^*] > 0$$

Multiplying both sides by the positive quantity $\frac{\sigma_{Pol}^4\sigma_{Her}^4\left(\frac{a_{ph}(t_p^*)H^*}{\sigma_{Her}^2} - \frac{a_{pm}(t_p^*)M^*}{\sigma_{Pol}^2}\right)^2}{[a_{pm}(t_p^*)M^*][a_{ph}(t_p^*)H^*](t_h - t_m)^2}$ preserves the equivalence:

$$Invasibility \Leftrightarrow [a_{ph}(t_p^*)H^* - a_{pm}(t_p^*)M^*] + \frac{\sigma_{Pol}^4\sigma_{Her}^4\left(\frac{a_{ph}(t_p^*)H^*}{\sigma_{Her}^2} - \frac{a_{pm}(t_p^*)M^*}{\sigma_{Pol}^2}\right)^3}{[a_{pm}(t_p^*)M^*][a_{ph}(t_p^*)H^*](t_h - t_m)^2} > 0$$

By defining $f(t_p^*)$ as:

| | |
|---|---|
| $f(t_p^*) = \sqrt{\frac{\sigma_{Pol}^4\sigma_{Her}^4\left(\frac{a_{ph}(t_p^*)H^*}{\sigma_{Her}^2} - \frac{a_{pm}(t_p^*)M^*}{\sigma_{Pol}^2}\right)^2}{[a_{pm}(t_p^*)M^*][a_{ph}(t_p^*)H^*](t_h - t_m)^2}}$ | $(A.6)$ |

We obtain:

$$Invasibility \Leftrightarrow \left[a_{ph}\left(t_p^*\right)H^* - a_{pm}\left(t_p^*\right)M^*\right] + f\left(t_p^*\right)^2 \left(\frac{a_{ph}\left(t_p^*\right)H^*}{\sigma_{Her}^2} - \frac{a_{pm}\left(t_p^*\right)M^*}{\sigma_{Pol}^2}\right) > 0$$

$$Invasibility \Leftrightarrow -a_{pm}\left(t_p^*\right)M^* \left[1 + \frac{f(t_p^*)^2}{\sigma_{Pol}^2}\right] + a_{ph}\left(t_p^*\right)H^* \left[1 + \frac{f(t_p^*)^2}{\sigma_{Her}^2}\right] > 0 \qquad (4)$$

Which corresponds to inequality (4) in the main document, thus ending the proof ■.

#### References


Dieckmann, U., and R. Law. 1996. The dynamical theory of coevolution: a derivation from stochastic ecological processes. Journal of Mathematical Biology 34:579–612.

Hutson, V., and R. Law. 1985. Permanent coexistence in general models of three interacting species. Journal of Mathematical Biology 21:285–298.

Sobol', I. Y. M. 1967. On the distribution of points in a cube and the approximate evaluation of integrals. USSR Computational Mathematics and Mathematical Physics 7:86–112.

Tomczak, M., and E. Tomczak. 2014. *The need to report effect size estimates revisited. An overview of some recommended measures of effect size*. Trends in Sport Sciences (Vol. 1).

Yacine, Y., and N. Loeuille. 2022. Stable coexistence in plant-pollinator-herbivore communities requires balanced mutualistic vs antagonistic interactions. Ecological Modelling 465:109857.

### Appendix B: Setting and exploring the parameter space

#### I. Choosing parameter values

The parameters that do not define plant-animal interactions are $(r_p, r_m, r_h, c_p, c_m, c_h, e_m, e_h)$. This set is referred to as the "ecological parameter set", and three such sets were studied (**Fig. B1.A**) chosen to cover vastly different scenarios in order to obtain a wide overview of potential eco-evolutionary dynamics. In contrast, parameters directly affecting plant-animal interactions – the "interspecific parameter set" $(t_h - t_m, a_{pm0}, a_{ph0}, \sigma_{Pol}, \sigma_{Her})$ – are at the core of our investigation. As such, we made them vary independently and systematically within interval ranges (**table 1**).

The values of our main ecological parameter set were chosen so that a wide range of plant-animal interaction strengths are compatible with stable coexistence (**Fig. B1.A.a**). The pollinator intrinsic growth rate is much higher than the herbivore one to favor positive population densities. Pollinators are, however, disfavored in terms of intraspecific competition and conversion efficiency to reduce the occurrence of unbounded population growths. Note that while unbounded growth may still happen (light blue area in **Fig. B1.A.a**), the plant-herbivore interaction considerably reduces the size of the associated region of parameter space, which would otherwise coincide with everything at the right of the vertical blue dotted line (**Fig. B1.A. a)**. In contrast to our main ecological parameter set, our second ecological parameter set (**Fig. B1.A.b**) does not favor any animal species over the other, and is thus referred to as "symmetrical". Finally, our third ecological parameter set makes unbounded population growth impossible over the range of variation of the interspecific parameter set (**Fig. B1.A.c**). This is achieved (Yacine and Loeuille 2022) by increasing plant and pollinator competition rates so that equation (B.1) is satisfied.

$$c_p c_m - e_m \{a_{pm}(t_p)\}_{max}^{\;2} > 0$$
$$\Leftrightarrow c_p c_m - e_m a_{pm}(t_p = t_m)^2 > 0$$
$$\Leftrightarrow c_p c_m - e_m \frac{a_{pm0}^{\;2}}{2\pi\sigma_{Pol}^{\;2}} > 0 \qquad (B.1)$$

It is sufficient for equation (B.1) to be satisfied for $a_{pm0} = 9$ and $\sigma_{Pol} = 1.5$ to guarantee it is satisfied over the whole range of variation of the intraspecific parameter set. With our third ecological parameter set, we obtain $c_p c_m - e_m \frac{a_{pm0}^{\;2}}{2\pi\sigma_{Pol}^{\;2}} = 1.7 \text{ x } 0.7 - 0.3 \text{ x } \frac{9^2}{\sqrt{2\pi} \text{ x } 1.5} \approx 0.044 > 0$. As a result, there is no region of plant phenotypic space $(t_p)$ associated with unbounded growth given this latter ecological parameter set, which is therefore labeled as "impossible unbounded growth".

Finally, the interval ranges over which interspecific parameters are allowed to vary (**table 1**) were chosen so that a wide variety of pollination and herbivory intensities are accessible (**Fig. B1.B**). Indeed, the vast majority of the stable coexistence area is covered when varying the interspecific parameters from one extreme configuration favoring pollination over herbivory (**Fig B1.B.a**), to the other extreme favoring herbivory over pollination (**Fig. B1.B.c**).

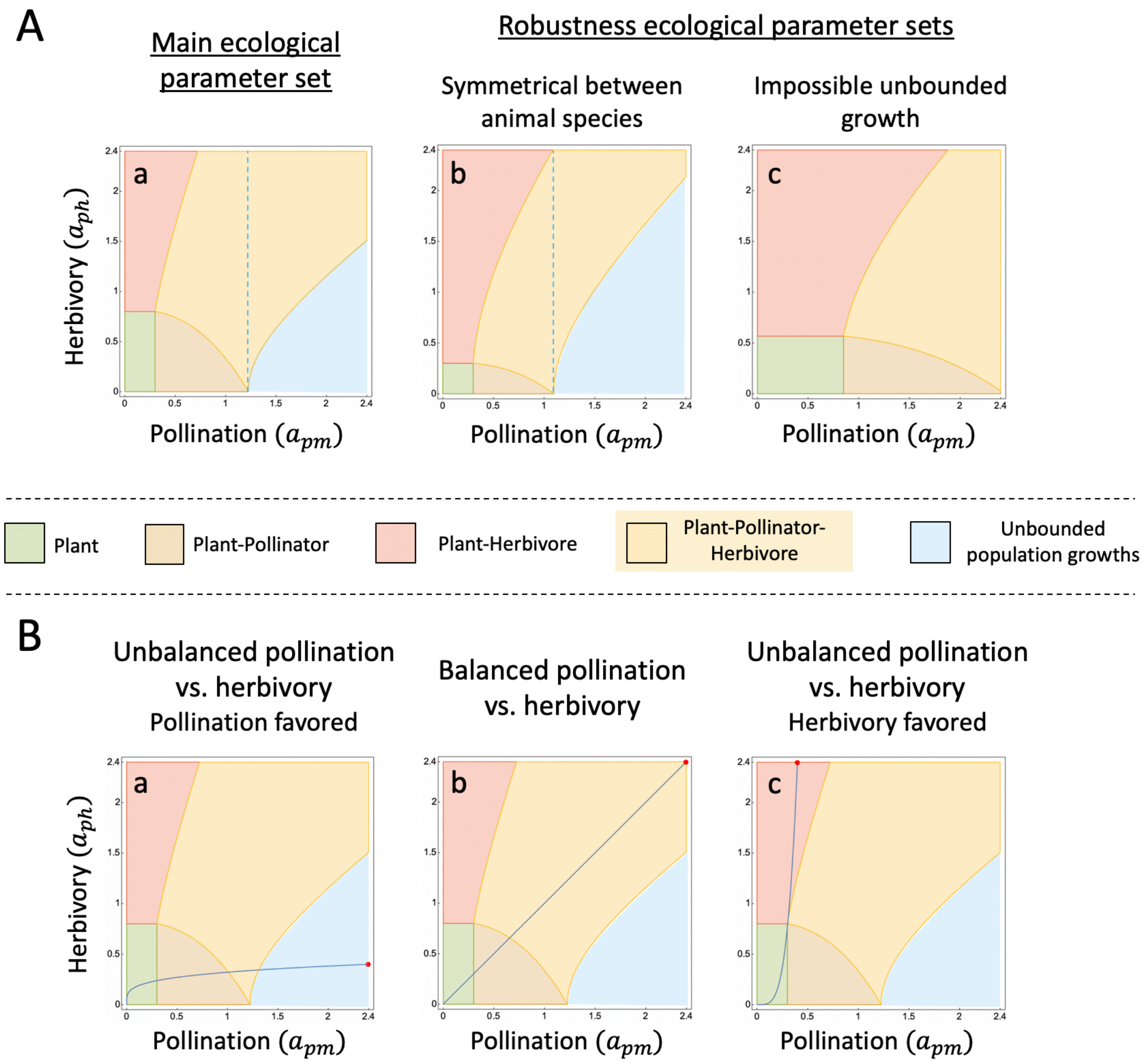


**Fig. B1: Community composition according to the strengths of pollination and herbivory. A. Choosing the ecological parameter sets. a.** The main ecological parameter set allows a wide stable coexistence area. $r_p = 10, r_m = -1, r_h = -4, c_p = 0.6, c_m = 0.5, c_h = 0.4, e_m = 0.2, e_h = 0.3$. **b. Robustness ecological parameter set** that is **symmetrical** between animal species. $r_p = 10, r_m = r_h = -1, c_p = 0.6, c_m = c_h = 0.4, e_m = e_h = 0.2$. **c. Robustness ecological parameter set** that makes **unbounded population growth impossible** over the ranges of interspecific parameters. $r_p = 10, r_m = r_h = -1, c_p = 1.7, c_m = 0.7, c_h = 0.4, e_m = 0.2, e_h = 0.3$. In **a & b**, the whole area at the right side of the blue dotted line would be associated with unbounded population growth in the absence of herbivores. **B. The range of the interspecific parameter set was chosen to allow a wide exploration of pollination and herbivory intensities. a.** Pollination is favored over herbivory ($t_h = t_m = 0, a_{pm0} = 9, a_{ph0} = 3, \sigma_{Pol} = 1.5, \sigma_{Her} = 3$). **b.** Balanced pollination vs. herbivory ($t_h = t_m = 0, a_{pm0} = 9, a_{ph0} = 9, \sigma_{Pol} = 1.5, \sigma_{Her} = 1.5$). **c.** Herbivory is favored over pollination ($t_h = t_m = 0, a_{pm0} = 3, a_{ph0} = 9, \sigma_{Pol} = 3, \sigma_{Her} = 1.5$). The blue curve corresponds to the sets of pollination and herbivory strengths accessible (i.e. $\left(a_{pm}(t_p), a_{ph}(t_p)\right)$ as the plant phenotype varies given the interspecific parameter set chosen (i.e. $(t_h - t_m, a_{pm0}, a_{ph0}, \sigma_{Pol}, \sigma_{Her})$). The red point indicates the position of $t_m$ and $t_h$. The ecological parameter set is the main one (Fig. S2A.a).

### II. Monte Carlo experiments

#### 1. Correspondence between evolutionary dynamics and types of selection.

In order to study how the interplay of plant-animal interactions affects the selection acting on the plant trait, we characterize the type of selection acting on each region of phenotypic space over which stable three-species coexistence is observed. As specified in the Method section, this is done by mapping the different types of selection – stabilizing, disruptive, runaway – into the evolutionary dynamics derived from the framework of adaptive dynamics. Using **Fig. B2**, we here illustrate how this mapping is implemented when no region of phenotypic space is characterized by unbounded population growth (**Fig. B2.b**), or when such an unbounded growth region boards the region of stable coexistence (**Fig. B2.c**).

<u>When unbounded growth is absent:</u>

1. Stabilizing selection occurs in the basin of attraction of a CSS. In **Fig. B3**, the proportion of phenotypic space corresponding to stabilizing selection is thus $\frac{Rep_1 - CSS}{t_{max} - t_{min}}$. Such quantities are summed if several CSSs occur.

2. Disruptive selection occurs in the basin of attraction of a BP. In **Fig. B3**, the proportion of phenotypic space corresponding to disruptive selection is thus $\frac{Rep_2 - Rep_1}{t_{max} - t_{min}}$. Such quantities are summed if several BPs occur.

3. Runaway selection occurs when neither stabilizing nor disruptive selection is occurring. This notably implies that selection is runaway when there are no evolutionary singularities (except when unbounded growth happens, see below). In **Fig. B3**, the proportion of phenotypic space corresponding to runaway selection is thus $\frac{t_{max} - Rep_2}{t_{max} - t_{min}}$. Such quantities are summed if runway selection occurs multiple times.

<u>When unbounded growth is present (additional rule to be applied before rule 3 just above):</u>

It is not straightforward to describe selection within unbounded growth regions as classical tools of adaptive dynamics, which require stable ecological equilibria, do not apply. The selective gradient is however defined in the vicinity of such regions and we found unbounded growth regions to be attractive in terms of evolution (i.e. convergence): as exemplified in **Fig. B2.c**, the selection gradient is positive in the phenotypic space below them (for $t_{min} < t_p <$

$t_{min}^{\infty}$), and negative above them (for $t_{max}^{\infty} < t_p < t_{max}$). This observation (it is a numerical result based on several tests, not an analytical result) motivates our choice to describe directional selection towards and within an unbounded growth region as stabilizing selection (please note this refers to evolutionary dynamics, and not ecological dynamics). Our choice is also coherent with the observation that in the E3 diagrams (x-axis $t_h - t_m$) where unbounded growth is observed **(Fig. 5b & 5d, Fig. S2d),** a CSS always collides with the area of unbounded growth, again as exemplified in **Fig. B2.c**. Our choice finally maintains the property of stabilizing selection being associated with the maintenance of coexistence. Indeed, areas of unbounded growth are regions of phenotypic space in which our model fails to produce realistic dynamics, but in which, from a biological point of view, coexistence should be maintained (notion of "permanent coexistence", Hutson and Law 1985, see discussion of Yacine and Loeuille 2022).

The calculation of the proportion of phenotypic space under each type of selection in the case of unbounded growth occurrence is illustrated in **Fig. B2.c**. There, the proportion of phenotypic space under stabilizing selection is $\frac{Rep_2 - t_{min}}{t_{max} - t_{min}}$, while that under runway selection is $\frac{t_{max} - Rep_2}{t_{max} - t_{min}}$. We thus consider the phenotypic region $[t_{min}^{\infty}, t_{max}^{\infty}]$to be under stabilizing selection.

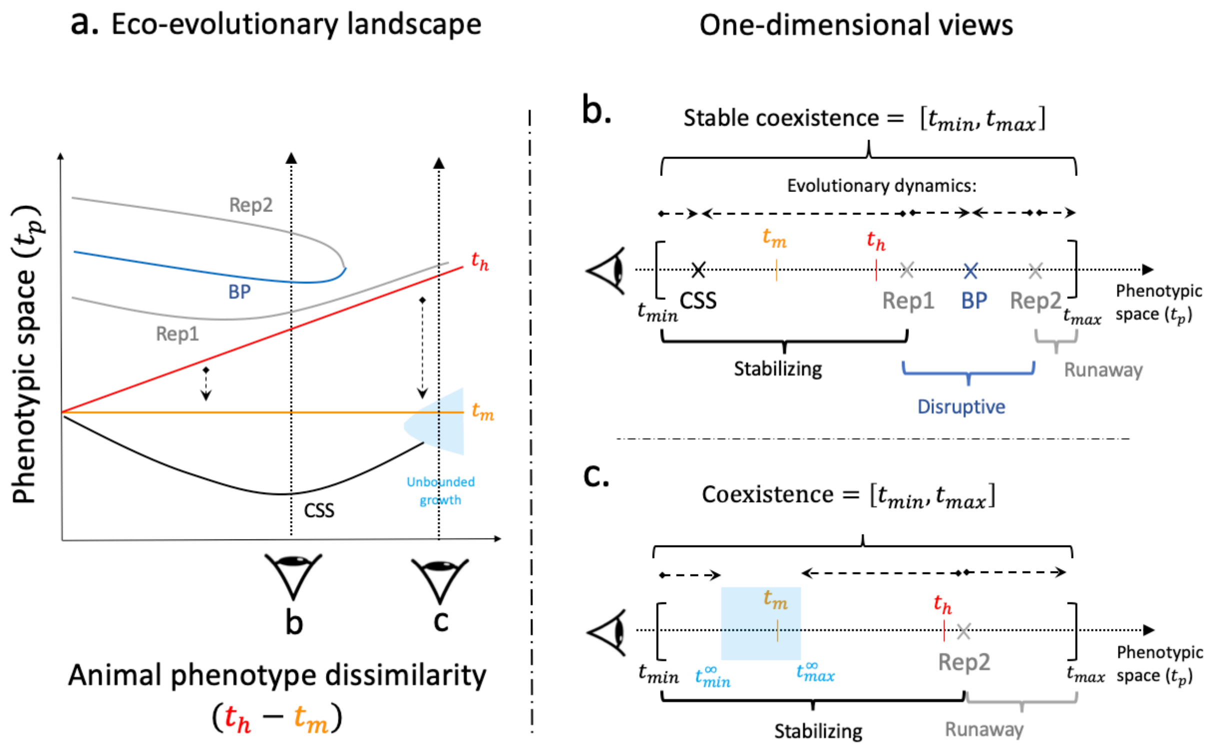


**Fig. B2: Correspondence between evolutionary dynamics and types of selection. a. Schematic eco-evolutionary landscape** characterized by one continuously stable strategy (CSS), one branching point (BP) and two repellors (Rep1 & Rep2). The graph corresponds to a schematic E3-diagram, while realistic ones are provided elsewhere, e.g. Fig. 2a or Fig. 5 in main document. **b-c. One-dimensional views of the eco-evolutionary landscape** illustrating how **the proportion of phenotypic space under each type of selection** is calculated in the **absence (b)** or **presence (c)** of unbounded growth regions (see text above).

#### 2. Relationship between the type of selection and the relative importance of pollination vs. herbivory (MC 1)

In a first Monte Carlo experiment (MC1), we quasi-randomly sample 10 000 interspecific parameter sets. The sampling corresponds to a 5-dimensional Sobol sequence (Sobol' 1967) which enables a low-discrepancy sampling (i.e. a more homogenous coverage of the parameter space). For each parameter set, we calculate the position and nature of singularities occurring in a stable coexistence context. We also calculate plant-animal interaction strengths and animal densities at each singularity. This allows linking the type of selection (locally) to the ecological context, i.e. the relative importance of pollination and herbivory (locally). Consistent with the correspondence between evolutionary dynamics and types of selection introduced in the previous section, we consider that a CSS corresponds to stabilizing selection, a BP corresponds to disruptive selection, and a repellor corresponds to runaway selection. Repellors leading to the disruption of stable plant-pollinator-herbivore coexistence are included (e.g. Rep2 in **Fig. B3**), while repellors surrounded by convergent singularities (e.g. Rep1 in **Fig. B3**) are excluded. Three ratios of pollination over herbivory are used to describe the ecological context: the ratio of interspecific interaction strengths at the singularity $t_p^*$ (i.e. $\frac{a_{pm}(t_p^*)}{a_{ph}(t_p^*)}$), the ratio of animal densities (i.e. $\frac{M^*}{H^*}$), and their more integrative product (i.e. $\frac{a_{pm}(t_p^*)M^*}{a_{ph}(t_p^*)H^*}$).

<u>Statistical analysis:</u>

For each ratio of pollination to herbivory, we tested its power to discriminate between the three types of selection – stabilizing, disruptive and runaway – using a Kruskal-Wallis test (non-parametric ANOVA, packages "rstatix" and "coin" in R). Based on Tomczak and Tomczak (2014), the effect size (i.e. proportion of variance explained) is computed as the eta squared based on the H-statistic (i.e.$\frac{H-k+1}{n-k}$, with $n$ and $k$ the total number of observations and the number of groups, respectively). We complemented the analysis by comparing the distribution of ratios between every two groups (i.e. every two types of selection) using the Wilcoxon-Mann-Whitney test (Holm-Bonferroni correction for multiple comparisons). Its effect size (Tomczak and Tomczak 2014) is computed as the Z-statistic divided by the square root of the total sample size $N$ (i.e. $\frac{Z}{\sqrt{N}}$). The tests were performed on the quasi-random (Sobol) draw of 10000 interspecific parameter sets, the ecological parameter set being fixed (focal one, see **Table 1**). Note that a given parameter set can lead to several types of selection (i.e. several evolutionary singularities), a dependence that we were not able to incorporate in the statistical model. The results of this statistical analysis are presented in **table S1** and **table C1** (**appendix C.I**). All statistical tests were highly significant, with $p_{value}$ always lower than $2.2\ 10^{-16}$.

#### 3. Effect of interspecific parameters on the selective landscape (MC 2)

Given a focal parameter from the interspecific parameter set (i.e. $t_h - t_m$, we divide its range (table 1) into six intervals of the same length. For each interval, we then sample 1000 interspecific parameter sets (5-dimensional Sobol sequence), but constrain the focal parameter within the considered interval. For each sampled parameter set, we calculate the proportion of phenotypic space corresponding to each type of selection (as in **Fig. B3**). The 1000 distributions obtained are combined (mean, standard error) to obtain a characterization of the selective landscape when the focal parameter is within the considered interval. The variation of the selective landscape while varying the constraining interval measures the effect of the focal interspecific parameter on selection. The results obtained from this Monte Carlo experiment are presented in **Fig. 4 & C2** in **appendix C.II**. Please note finally that animal phenotype dissimilarity $(t_h - t_m)$ was constrained within $[0, 1.5]$ (instead of $[0, 3]$) when the focal interspecific parameter was among the other four (i.e. $a_{pm0}, a_{ph0}, \sigma_{Pol}, \sigma_{Her}$). This was done to better capture the effects of these parameters on disruptive selection, the latter being much less frequent when animal phenotype dissimilarity is high (i.e. for weak trade-offs, see Fig. 4A and main text).

#### References


Hutson, V. & Law, R. (1985). Permanent coexistence in general models of three interacting species. *J. Math. Biol.*, 21, 285–298.

R Core Team (2022). R: A language and environment for statistical computing. R Foundation for Statistical Computing, Vienna, Austria. URL https://www.R-project.org/.

Sobol', I. Y. M. 1967. On the distribution of points in a cube and the approximate evaluation of integrals. USSR Computational Mathematics and Mathematical Physics 7:86–112.

Tomczak, M., and E. Tomczak. 2014. *The need to report effect size estimates revisited. An overview of some recommended measures of effect size.* Trends in Sport Sciences (Vol. 1).

Yacine, Y., and N. Loeuille. 2022. Stable coexistence in plant-pollinator-herbivore communities requires balanced mutualistic vs antagonistic interactions. Ecological Modelling 465:109857.

### Appendix C: Robustness to the variation of the ecological parameter set

#### I. Relationship between type of selection and relative strength of pollination vs. herbivory (MC1)

We investigate the relationship between selection type and pollination to herbivory ratio in the case of our two robustness ecological parameter sets (values in parenthesis in **table 1**).

The results presented in **Fig. 3** are robust to the variation of the ecological parameter set (**Fig. C1**):

- Stabilizing selection is characterized by large pollination to herbivory ratios, while runaway selection is characterized by small pollination to herbivory ratios.
- Disruptive selection is characterized by balanced pollination to herbivory ratios, in between the ratios favoring the two other types of selection
- The ratio of pollination to herbivory largely explains the nature of selection (Kruskal-Wallis effect size of 71% & 67%).

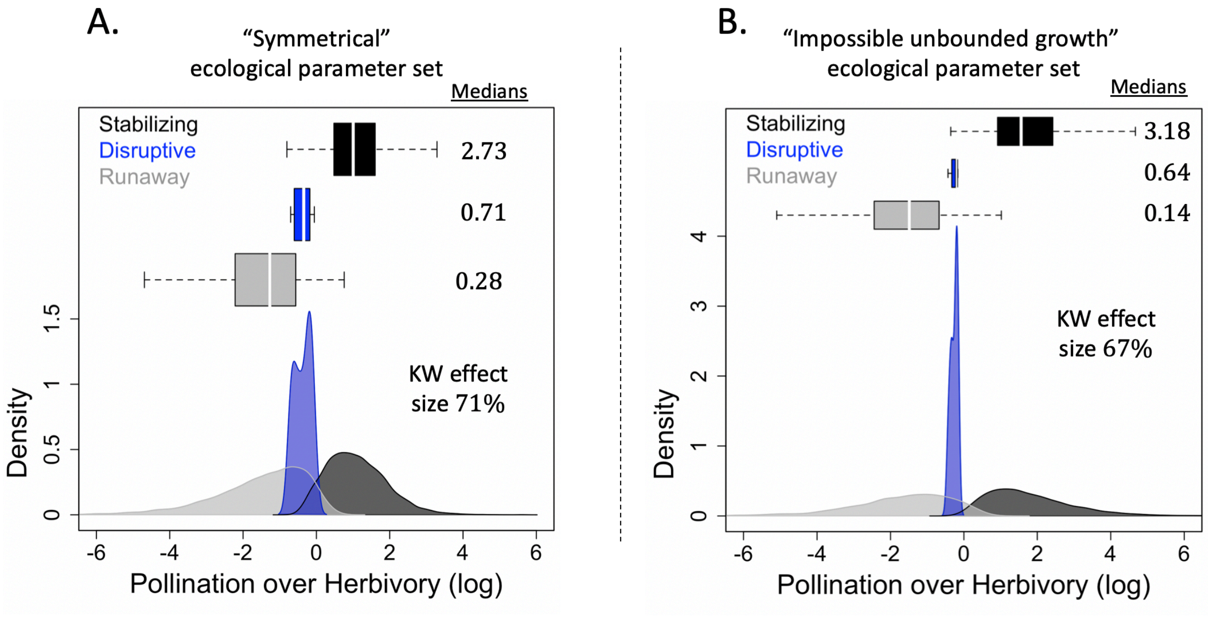


$$\left.\frac{a_{pm}M^*}{a_{ph}H^*}\right]_{t_p^*}$$

**Fig. C1: Distribution of pollination to herbivory ratios depending on selection type for our robustness ecological parameter sets (A. Symmetrical, B. Impossible unbounded growth).** The ratio is measured at the corresponding evolutionary singularity. The medians indicated correspond to absolute values (no log). The effect size of the Kruskal-Wallis test ($p_{value} < 2.2\ 10^{-16}$) is also indicated. Ecological parameter set: ($r_p = 10, r_m = -1, r_h = -1, c_p = 0.6\ (A)\ or\ 1.7\ (B), c_m = 0.4\ (A)\ or\ 0.7\ (B), c_h = 0.4, e_m = 0.2, e_h = 0.2\ (A)\ or\ 0.3\ (B)$). Results are from our first Monte Carlo experiment (MC1, appendix B.II.2)

Results of the statistical analysis

Overall, the variation of the ecological parameter does not change the results derived so far (Table S1). Indeed, the three ratios of pollination to herbivory still largely explain the nature of selection (Kruskal-Wallis effect size). An important difference is that disruptive selection is much less frequent, with 20 (resp. 11) branching points encountered across the 10 000 sampled interspecific parameter sets in the case of our "symmetrical" ecological parameter set (resp. "impossible unbounded growth" ecological parameter set.)

| Ecological parameter set | Ratio of pollination to herbivory | Expression | Median per type of selection | | | Kruskal-Wallis effect size |
|---|---|---|---|---|---|---|
| | | | Runaway | Disruptive | Stabilizing | |
| Symmetrical between animal species | Interaction strengths and animal densities | $\left.\frac{a_{pm}M^*}{a_{ph}H^*}\right]_{t_p^*}$ | 0.24 | 1.23 | 3.10 | 0.713 |
| | Interaction strengths | $\left.\frac{a_{pm}}{a_{ph}}\right]_{t_p^*}$ | 0.66 | 1.04 | 1.52 | 0.750 |
| | Animal densities | $\left.\frac{M^*}{H^*}\right]_{t_p^*}$ | 0.36 | 1.15 | 2.01 | 0.750 |
| Impossible unbounded population growth | Interaction strengths and animal densities | $\left.\frac{a_{pm}M^*}{a_{ph}H^*}\right]_{t_p^*}$ | 0.18 | 1.38 | 4.32 | 0.678 |
| | Interaction strengths | $\left.\frac{a_{pm}}{a_{ph}}\right]_{t_p^*}$ | 1.18 | 1.57 | 1.96 | 0.677 |
| | Animal densities | $\left.\frac{M^*}{H^*}\right]_{t_p^*}$ | 0.16 | 0.89 | 2.07 | 0.678 |

**Table C1: Results of the statistical analysis for the three ratios of pollination to herbivory tested in the case of the robustness ecological parameter sets.** Sample size (interspecific parameter sets): 10000, which resulted in 7031 Stabilizing, 7301 Runaway and 20 Disruptive (Symmetrical); and 4214 Stabilizing, 2214 Runaway and 11 Disruptive (Impossible unbounded growth). Ecological parameter sets provided in Fig. C1 above. All statistical tests were highly significant, with a $p_{value}$ below $2.2\ 10^{-16}$ .

#### II. Effect of interspecific parameters on the selective landscape (MC 2)

Here, we present (**Fig. C2 & C3**) the effect of varying the parameters $t_h - t_m, a_{pm0}, a_{ph0}, \sigma_{Pol}$ or $\sigma_{Her}$ on the selective landscape in the case of our two robustness ecological parameter sets, the symmetrical one (**Fig. C2**) and the one in which unbounded growth is not possible (**Fig. C3**).

The effect of trade-off strength on selection varies with the ecological parameter set. For the “symmetrical” ecological parameter set (**Fig. C2.A**), decreasing trade-off strength is responsible for an increase (resp. decrease) in the prevalence of stabilizing (resp. runaway) selection. The opposite is observed under our main ecological parameter set (**Fig. 4A**). For the “impossible unbounded growth” parameter set (**Fig. C3.A**), the relationships are non-monotonical: the prevalence of stabilizing (resp. runaway) selection is maximized (resp. minimized) at intermediate trade-offs. The effect of trade-off on the prevalence of disruptive selection is however robust as such a type of selection is only possible for strong and intermediate trade-offs, but not when the trade-off is weak (**Fig. C2.A & C3.A**).

The effects of basal interaction rates on selection seem robust to the variation of the ecological parameter set. An increase in the basal pollination rate ($a_{pm0}$) makes stabilizing selection more frequent at the expense of runaway selection (**Fig C2.B.a & C3.B.a**). The effect of the basal herbivory rate ($a_{ph0}$) is utterly opposite (**Fig. C2.B.b & C3.B.b**). In contrast, the effects of interaction niche widths on selection are not robust to the variation of the ecological parameter set. For the “symmetrical” parameter set, the niche width of both interactions seems to similarly affect selection: an intermediate niche width maximizes runaway selection and minimizes stabilizing selection (**Fig. C2.B. c & d**). For the “impossible unbounded growth” ecological parameter set, a similar relationship is found for herbivory niche width (**Fig. C3.d**), while pollination niche width has almost no effect on the relative prevalence of selection types. Note finally that disruptive selection is extremely rare for the robustness ecological parameter sets (**Fig. C2 & C3,** see also **table D1** in **appendix D**).

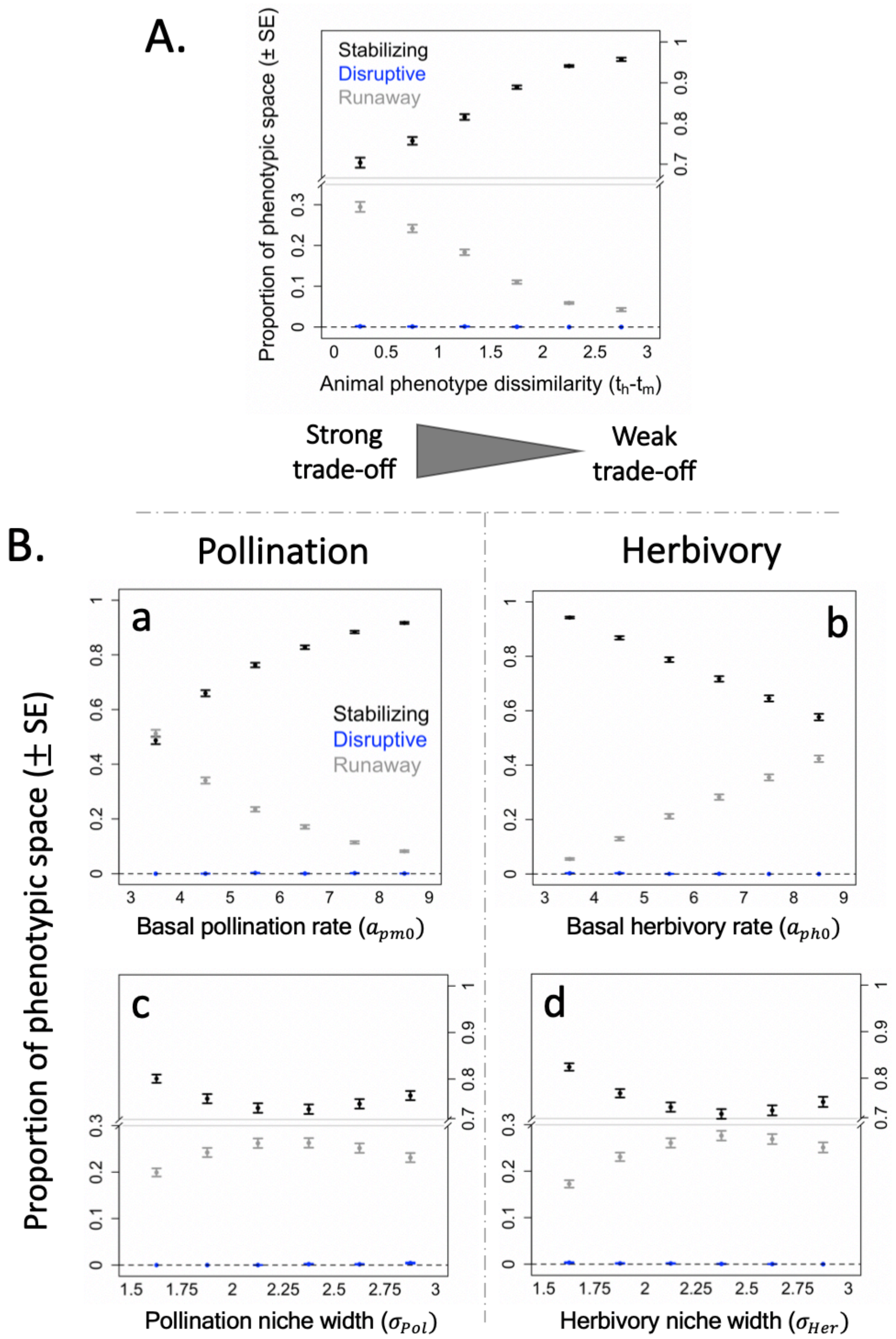


**Fig. C2: Effect of interspecific parameters on the selective landscape for our "symmetrical" robustness ecological parameter set. A. The selective landscape depends on trade-off intensity. B. Opposite effects of pollination and herbivory on selection.** Please note that animal phenotype dissimilarity $(t_h - t_m)$ was here further constrained within $[0, 1.5]$. **a. b. Variations in basal interaction rates. c. d. Variations in niche widths.** Results are from our second Monte Carlo experiment (MC 2, appendix B.II.3), with 1000 interspecific parameter sets sampled at each point. Y-axes indicate the normalized size of the basins of attraction (Mean $\pm$ SE) associated with each type of selection (see **appendix B.II.1**). Ecological parameter set: $(r_p = 10, r_m = -1, r_h = -1, c_p = 0.6, c_m = c_h = 0.4, e_m = 0.2, e_h = 0.2)$.

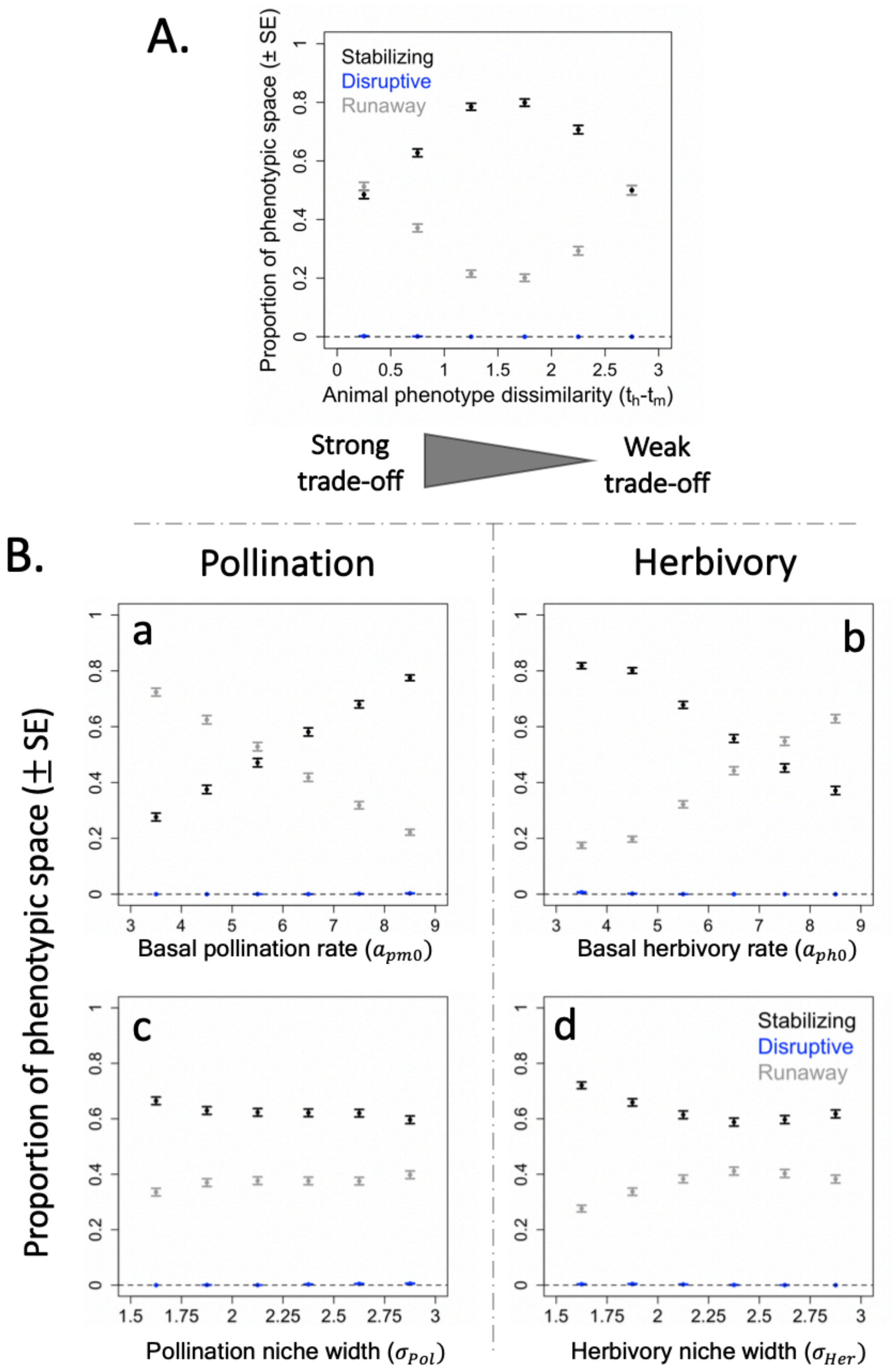


**Fig. C3: Effect of interspecific parameters on the selective landscape for our "impossible unbounded growth" robustness ecological parameter set. A. The selective landscape depends on trade-off intensity. B. Opposite effects of pollination and herbivory on selection.** Please note that animal phenotype dissimilarity $(t_h - t_m)$ was here further constrained within $[0, 1.5]$. **a. b. Variations in basal interaction rates. c. d. Variations in niche widths.** Results are from our second Monte Carlo experiment (MC 2, appendix B.II.3), with 1000 interspecific parameter sets sampled at each point. Y-axes indicate the normalized size of the basins of attraction (Mean ± SE) associated with each type of selection (see **appendix B.II.1**). Ecological parameter set: $(r_p = 10, r_m = -1, r_h = -1, c_p = 1.7, c_m = 0.7, c_h = 0.4, e_m = 0.2, e_h = 0.3)$.

### Appendix D: Joint selection and the emergence of plant dimorphism

In order to assess the maintenance of plant dimorphism following branching points, we simulated the eco-evolutionary dynamics associated with all the branching points encountered in our second Monte Carlo experiment (MC2, appendix B.III.3), when constraining the sampling of trade-off intensity (6 x 1000 samplings of interspecific parameter sets). 676 branching points were encountered in the case of our main ecological parameter set, 15 in the case of our second ("symmetrical") ecological parameter set, and 7 in the case of our third ecological parameter set ("impossible unbounded growth") . The number and proportion of maintained dimorphism observed at each range of trade-off intensity is given in **table D1**. In all cases where dimorphism was not maintained, evolution ended at a CSS at which stabilizing selection prevailed. Note that we were not able to assess the maintenance of dimorphism in the case of one branching point occurring at intermediate trade-off ($1 < t_h - t_m \leq 1.5$, main ecological parameter set). Disruptive selection was weak, so that a large number of morphs were accumulating at the BP-phenotype, making the simulation extremely slow: the emergence of two branches was not observed after 5 days of simulations. To be conservative, we considered that dimorphism was not maintained in that case.

| Ecological Parameter set | Ecological trade-off | | Branching Points | Dimorphism maintenance | **Relative frequency of dimorphism maintenance** |
|---|---|---|---|---|---|
| Main ($r_p = 10, r_m = -1, r_h = -4, c_p = 0.6, c_m = 0.5, c_h = 0.4, e_m = 0.2, e_h = 0.3$) | Strong | $0 < t_h - t_m \leq 0.5$ | 155 | 41 | 26.5% |
| | | $0.5 < t_h - t_m \leq 1$ | 160 | 0 | 0% |
| | Intermediate | $1 < t_h - t_m \leq 1.5$ | 169 | 0 | 0% |
| | | $1.5 < t_h - t_m \leq 2$ | 108 | 0 | 0% |
| | Weak | $2 < t_h - t_m \leq 2.5$ | 61 | 0 | 0% |
| | | $2.5 < t_h - t_m \leq 3$ | 23 | 0 | 0% |
| | All | $0 < t_h - t_m \leq 3$ | 676 | 41 | 6% |
| Symmetrical ($r_p = 10, r_m = r_h = -1, c_p = 0.6, c_m = c_h = 0.4, e_m = e_h = 0.2$) | Strong | $0 < t_h - t_m \leq 0.5$ | 4 | 0 | 0% |
| | | $0.5 < t_h - t_m \leq 1$ | 5 | 0 | 0% |
| | Intermediate | $1 < t_h - t_m \leq 1.5$ | 3 | 0 | 0% |
| | | $1.5 < t_h - t_m \leq 2$ | 3 | 0 | 0% |
| | Weak | $2 < t_h - t_m \leq 3$ | 0 | 0 | 0% |
| | All | $0 < t_h - t_m \leq 3$ | **15** | **0** | 0% |

| | | | | | |
|---|---|---|---|---|---|
| Impossible unbounded growth ($r_p = 10, r_m = r_h = -1, c_p = 1.7, c_m = 0.7, c_h = 0.4, e_m = 0.2, e_h = 0.3$) | Strong | $0 < t_h - t_m \leq 0.5$ | 5 | 0 | 0% |
| | | $0.5 < t_h - t_m \leq 1$ | 2 | 0 | 0% |
| | Intermediate and weak | $1 < t_h - t_m \leq 3$ | 0 | 0 | 0% |
| | **All** | $0 < t_h - t_m \leq 3$ | **7** | 0 | 0% |

**Table D1: Quantifying the maintenance of dimorphism as a result of disruptive selection.**
